# *Mycobacterium ulcerans* challenge strain selection for a Buruli ulcer controlled human infection model

**DOI:** 10.1101/2024.02.08.579445

**Authors:** Stephen Muhi, Andrew H. Buultjens, Jessica L. Porter, Julia L. Marshall, Marcel Doerflinger, Sacha J. Pidot, Daniel O’Brien, Paul D. R. Johnson, Caroline Lavender, Maria Globan, James McCarthy, Joshua Osowicki, Timothy P. Stinear

## Abstract

Critical scientific questions remain regarding infection with *Mycobacterium ulcerans*, the organism responsible for the neglected tropical disease, Buruli ulcer (BU). A controlled human infection model has the potential to accelerate our knowledge of the immunological correlates of disease, to test prophylactic interventions and novel therapeutics. Here we present microbiological evidence supporting *M. ulcerans* JKD8049 as a suitable human challenge strain. This non-genetically modified Australian isolate is susceptible to clinically relevant antibiotics, can be cultured in animal-free and surfactant-free media, can be enumerated for precise dosing, and has stable viability following cryopreservation. Infectious challenge of humans with JKD8049 is anticipated to imitate natural infection, as *M. ulcerans* JKD8049 is genetically stable following *in vitro* passage and produces the key virulence factor, mycolactone. Also reported are considerations for the manufacture, storage, and administration of *M. ulcerans* JKD8049 for controlled human infection.

## Introduction

The neglected tropical disease, Buruli ulcer (BU), is a bacterial infection of subcutaneous tissue caused by *Mycobacterium ulcerans*. A controlled human infection model (CHIM) has the potential to revolutionise our understanding of the immunological correlates of disease. By randomising participants to either an experimental or control arm, such a model would also enable researchers to rapidly test prophylactic and therapeutic interventions, development of which may be supported by studying immunobiological responses to experimental infection. We recently introduced 10 guiding selection criteria for an ideal *M. ulcerans* challenge strain [1]. Here, we describe our work to produce a standardised, low-dose inoculum suitable for strain manufacture, storage, pre-clinical testing and administration. Previously, we introduced *M. ulcerans* JKD8049 as a potential candidate challenge strain, matching a number of our guiding criteria [1]. JKD8049 was isolated in 2004 from a middle-aged Caucasian male in Point Lonsdale, Victoria, Australia, who presented with a non-severe painless ulcer on the posterior calf, the typical clinical BU syndrome seen in south-eastern Australia. This isolate is amenable to challenge by a biologically plausible route, as observed in a murine subcutaneous puncture model [2].

Here, we report investigation of *M. ulcerans* JKD8049 as a candidate human challenge strain against the 7 remaining aforementioned criteria [1], with comparisons to a panel of geographically diverse *M. ulcerans* isolates. Namely, we aimed to:

1. Demonstrate susceptibility to clinically relevant antibiotics,
2. Culture isolates in a non-toxic, animal-free medium without genetic or chemical modification,
3. Accurately enumerate bacilli to ensure consistent dosing,
4. Demonstrate viability after cryopreservation and thawing,
5. Confirm production of mycolactone *in vitro,*
6. Display a conserved repertoire of genes encoding candidate vaccine antigens,
7. Demonstrate genetic stability following serial passage.

We demonstrate that JKD8049 fulfils these remaining criteria and is a suitable candidate for further characterisation in a mammalian subcutaneous infection model. Our findings will guide the manufacture of *M. ulceran*s challenge doses, in accordance with the principles of Good Manufacturing Practice.

## Materials and methods

### Bacterial strains

A range of geographically diverse clinical isolates were selected to compare antibiotic susceptibility and growth characteristics (Table 1). Although there are numerous *M. ulcerans* isolates available for consideration, *M. ulcerans* isolates are remarkably conserved, with little genetic diversity [3]. The analysed *M. ulcerans* strains did not include genetically modified organisms. A dedicated working stock was cryopreserved to avoid external manipulation or contamination.

**Table 1:** Strains used in this study.

| Strain | Origin | Year isolated | SNP difference from JKD8049 | Mycolactones | Notes |
| --- | --- | --- | --- | --- | --- |
| **JKD8094**<br>ITM 97-1116 | Benin | 1997 | 3,691 | A/B | Clinical isolate, plaque [4] |
| **NM20/02** | Ghana | 2001-2003 | 3,659 | A/B | Passaged, attenuated [5] |
| **JKD8049** | Victoria, Australia | 2004 | *Ref.* | C* | Clinical isolate, ulcer [6,7] |
| **JKD8095**<br>ITM 98-0912 | China | 1998 | 21,990 | D<sup>7</sup> | Clinical isolate, ulcer [8] |
| **JKD8097**<br>ITM 06-3844 | Belgium (fish) | 2008 | 20,495 | F<sup>10</sup> | Unable to grow above 30°C [9] |

\*Known to produce a fraction of mycolactone A/B

### Whole genome sequencing

Whole genome sequencing (Illumina, San Diego, CA, USA) was used to confirm the isolate identities and their phylogenetic relationships (Table 1, Fig. 3). Snippy v4.4.5 was used to map read sets and assembled contigs against the fully assembled JKD8049 chromosome (GenBank accession: CP085200.1) using a ‘minfrac’ setting of 0.8 (https://github.com/tseemann/snippy). Comparisons of genomes were conducted using the snippy-core command with repetitive regions masked to mitigate the risk of false-positive variant calls and unreliable mapping. A maximum likelihood phylogenomic tree was built with FastTree v2.1.10 [10] using the core single nucleotide polymorphism (SNP) alignment produced by snippy-core.

### Antibiotic susceptibility testing

To determine the minimum inhibitory concentration (MIC) of each selected strain to rifampicin, clarithromycin and ciprofloxacin (Fig. 1), mycobacteria were cultured on Middlebrook 7H10 agar (BD, Sparks, MD, USA) enriched with 10% oleic acid, albumin, dextrose, and catalase (OADC; HiMedia, Mumbai, India). After 6 to 12 weeks of incubation (depending on the growth characteristics of each strain) at 30°C, culture material was scraped and transferred by 10 μL sterile loop into 13 mL flat-bottomed centrifuge tubes containing 10 sterilised 3 mm glass beads and 500 μl sterile phosphate buffered saline (PBS); the low initial volume enabled greater contact with the glass beads to optimise de-clumping. This tube was vortexed at high speed for 3 minutes, then a further 3.5 mL of PBS was added and vortexed again for 60 seconds. The tube was allowed to stand for 15 minutes, for large bacterial clumps to settle under gravity. The bacterial density was measured using a nephelometer (DensiCheck, Biomerieux, Durham, USA). Carefully avoiding settled clumps, the supernatant was then transferred by P1000 micropipette into a separate 50 mL Falcon tube, where it was adjusted to 0.5 MacFarland standard using sterile PBS. The suspension was then diluted 1:5 using sterile PBS and vortexed briefly. A 500 μl volume of this preparation was used to inoculate duplicate Mycobacteria Growth Indicator Tubes (MGITs; BD, Sparks, MD, USA) containing 4 mL 7H9 broth and 500 μl OADC containing doubling dilutions of rifampicin, clarithromycin or ciprofloxacin (Sigma-Aldrich, St. Louis, MO, USA) starting at 8 μg/mL. As per the MGIT procedure manual [11], the bacterial suspension was further diluted 1:100, and 500 μl was added to an antibiotic-free growth control tube containing diluent in OADC (dimethyl sulfoxide, 0.1M hydrogen chloride or acetone for rifampicin, ciprofloxacin and clarithromycin, respectively). Growth control tubes without further dilution were also included.

**Fig. 1.**
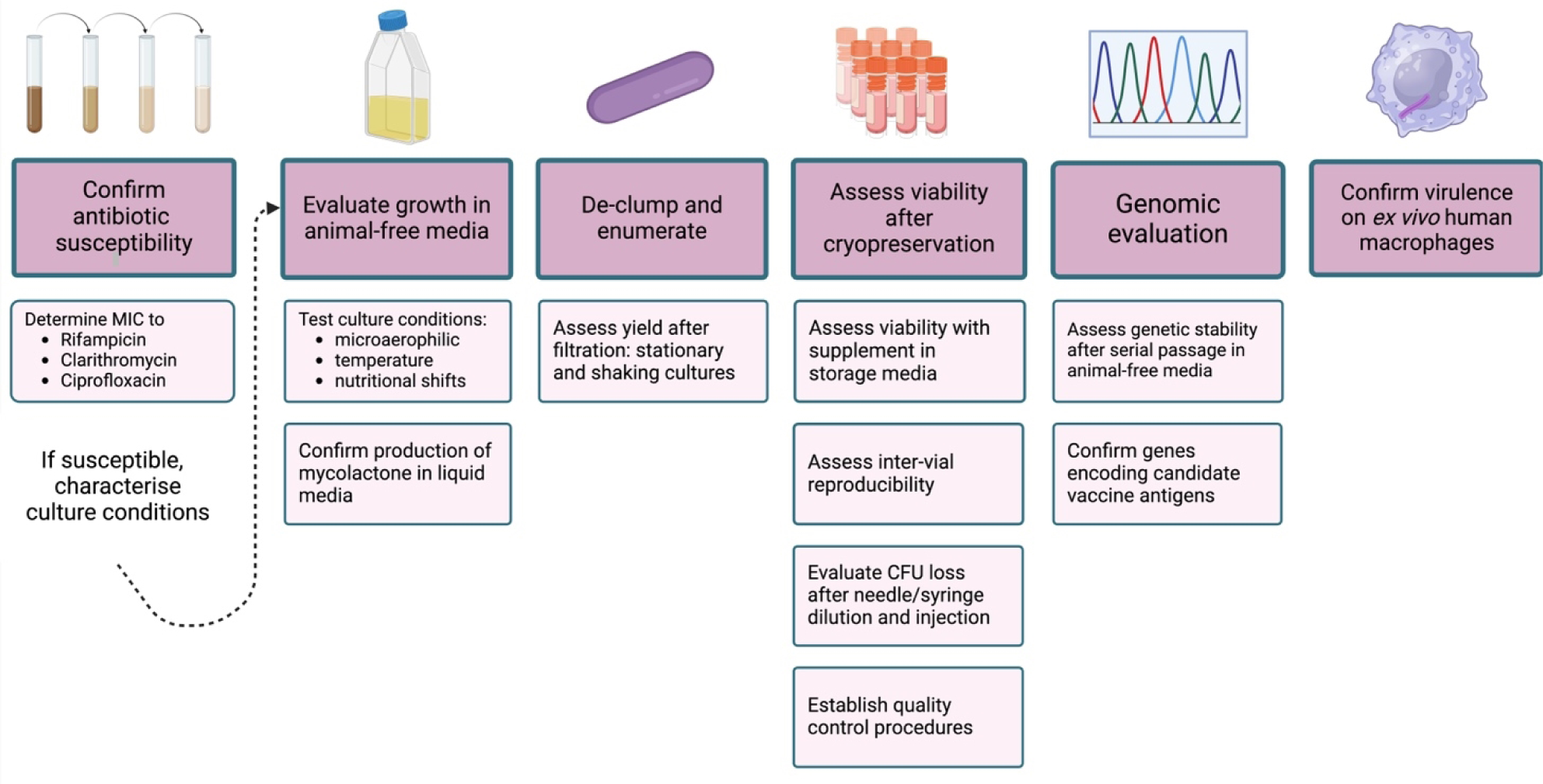
Experimental approaches for evaluating candidate *M. ulcerans* isolates for use in a CHIM. MIC: minimum inhibitory concentration.

Following incubation at 30°C, samples were reviewed twice daily for two weeks, then daily thereafter by detecting fluorescence using a manual MGIT reader (BD BACTEC MicroMGIT, USA). MIC determination was adapted from recommended methodology [11], such that when the growth control tubes demonstrated fluorescence, the antibiotic-containing tubes with the lowest drug-concentration displaying no fluorescence were considered to contain the MIC. The MGIT reader was calibrated in accordance with the manufacturer’s instructions for use (IFU). Antibiotics in solvent (at 1 μg/mL) without bacteria were used as negative controls. For quality control, 10 μL from each positive growth control tube was streaked onto non-selective nutrient agar and incubated at 37°C. Growth control tubes were also confirmed to contain acid-fast bacilli using Ziehl-Neelsen staining. MGITs were only opened briefly to add media or bacteria, in order to minimise oxygen loss.

A quality control isolate, *M. fortuitum* ATCC6841, was used to validate the performance of this methodology, as suggested in the MGIT procedure manual [11]. The procedure was performed identically to the test *M. ulcerans* isolates, although the growth control tube was diluted 1:5000 rather than 1:100 [11]. This isolate has published MIC values for rifampicin, with MIC ≥ 64 μg/mL reported in online databases (http://antibiotics.toku-e.com, accessed 7 February 2024) and from other reported methodologies [12], and MIC 16 μg/mL by agar dilution [13]. Reported clarithromycin MICs are variable, with values of 2 μg/mL reported by agar dilution [13], although reported ranges between 0.125 to 8 μg/mL have lead authors to conclude that the variability of susceptibility makes clarithromycin a less reliable quality control antibiotic for this isolate [14,15]. The isolate has a reportedly low MIC to ciprofloxacin, with MIC of 0.064 μg/mL by agar dilution [13].

### Culture conditions for manufacture of candidate challenge strains

Sauton’s media was prepared using an established recipe [16]. Ingredients for 1 L of liquid media (2 g citric acid, 0.5 g KH_2_PO4, 0.5 g MgSO_4_.7H_2_O, 0.05 g ferric ammonium citrate, and 4 g asparagine) were dissolved in 300 mL of distilled water and stirred with a magnetic stirring bar for 20 minutes, after which 60 mL of glycerol (of plant origin) and distilled water were added to a final volume of 1 L. Liquid media containing an animal-free and non-genetically modified growth supplement were made by adding 3.1 g of vegetable peptone broth (Veggietones VG0101, Oxoid, Basingstoke, UK) to 100 mL of distilled water (as per IFU) and stirred for 20 minutes until fully dissolved. This was then added to 900 mL of Sauton’s media (10% vol/vol). While stirring, 10 M NaOH was used to adjust the final pH to 7.4 prior to autoclaving at 121°C for 15 minutes, then stored at room temperature. This liquid media has a clear, light yellow colour, abbreviated ‘SMVT’ (Sauton’s media with ‘Veggietones’).

### Optimising culture conditions for homogeneous growth

Stationary cultures in SMVT were created by inoculating ≥ 1 McFarland of *M. ulcerans* suspension into five sealed tissue culture flasks each containing 50 mL of SMVT. Stationary cultures were then incubated at 30°C in standard atmospheric conditions. To remove clumps, the methodology described by Cheng and colleagues [17] was adapted. At two-weekly intervals, one stationary liquid culture was poured into a 50 mL syringe and plunged through a sterile 5 μm membrane filter (Millex-SV, Merck, Darmstadt, Germany), with slow and consistent pressure, until all material had passed through the filter. The filtrate was collected in a sterile container and enumerated via spot plating using six 3 μL replicates per plate, in serial dilution, on 7H10/OADC Middlebrook agar. Plates were incubated at 30°C and checked fortnightly for up to 12 weeks. Spread plates were performed to compare colony morphology before and after filtration (Fig. 4).

Orbital shaking cultures were established using 30 mL universal containers with 20 – 25 sterile 3 mm diameter glass beads and 6 mL SMVT. The media was inoculated by transferring culture material directly from an agar surface using a sterile 10 μL loop (as colonies are rough and waxy, the material was scraped or lifted off the agar surface and carefully transferred). Inoculated containers were set to shake at 200 rpm at 30°C. Cling film (Parafilm-M, Amcor, Zürich, Switzerland) was used to secure the screw-top lid on universal containers to minimise evaporative water loss and to further minimise contamination, considering the long shaking incubation period. Before harvesting, shaking was paused for 15 – 20 minutes for large clumps to settle with gravity. Drawing from the centre of the liquid culture, avoiding the surface pellicle or the settled culture material, samples were pipetted by P1000 micropipette into a 10 mL syringe with the plunger removed. The liquid was then slowly plunged through a 5 μm membrane filter, and the filtrate collected and swirled by hand for 10 seconds. Volumes removed for colony forming unit (CFU) enumeration were replaced with the original culture media (maximum volume 50 μL per timepoint) to avoid concentrating bacteria over time. Enumeration of bacilli, both pre- and post-filtration, was performed using 5 μL spots, with six technical replicates per plate, at serial dilution (10^-1^ to 10^-5^) on 7H10/OADC agar. Agar plates were incubated at 30°C and reviewed fortnightly. CFUs were manually counted, selecting the dilution containing between 3 and 30 CFU per spot. For comparison, triplicate cultures in Sauton’s medium and in Middlebrook 7H9 with albumin, dextrose and catalase (ADC) were established using the same methodology. Serial dilutions were created using the liquid medium being tested in each respective experiment (i.e., Sauton’s media, SMVT or 7H9/ADC). Negative control shaking cultures were also established at the time of inoculation and incubated for the length of each experiment. Ziehl-Neelsen stains before and after filtration were performed using 20 μL of each filtrate from all isolates, air dried, heat-fixed and viewed at 100x magnification with oil immersion. At least 30 high-power fields of the filtrate sample were inspected to confirm that no clumps were visible in the filtrate.

### Culture conditions

Microaerophilic conditions were established using CampyGen sachets (Thermo Fisher, Landsmeer, the Netherlands) placed within a sealed container, alongside a Yeast Casitone Fatty Acids Agar (YCFA) plate as an indicator (appearing pink in the presence of atmospheric oxygen). Continuous temperature monitoring of isolates cultured at both 30°C and 37°C was performed using a portable electronic thermometer in each incubator (Tinytag View 2, Gemini Data Loggers, UK).

### Viability following cryopreservation

Glycerol storage medium was produced by adding 20 mL glycerol to 80 mL of distilled water (20% vol/vol), autoclaved at 121°C for 15 minutes and stored at room temperature. Immediately following filtration, 0.25 mL of each isolate’s filtrate was transferred by pipette into a sterile 2 mL sterile screw-capped cryovial containing 0.5 mL of glycerol storage medium (i.e., 1:3 dilution). Samples were vortexed for 10 seconds and frozen promptly at -80°C. As the local standard cryopreservative contains tryptone soya broth in addition to glycerol, samples of JKD8049 (prepared as shaking cultures and filtered, as described earlier) were tested using Veggietones (Oxoid, ThermoFisher Scientific) in glycerol, with Veggietones replacing the tryptone soya broth at the same concentration. Glycerol (20% v/v) without any supplemental agent was used as a control. Veggietones glycerol storage media was prepared by adding 1.6 g of Veggietones powder to 80 mL of distilled water and stirring for 15 minutes. This was added to 20 mL glycerol, autoclaving for 15 minutes at 121°C, and stored at room temperature.

### Quality control

After thawing cryovials, quality control testing was performed by inoculating duplicate (1) nutrient agar (incubated for 4 weeks at 37°C) for general environmental contaminants, (2) brain-heart infusion broth (incubated for 4 weeks at 37°C) for fastidious contaminants, (3) Sabouraud agar (incubated for 6 weeks at 30°C) for fungal contaminants and (4) 7H10/OADC agar (incubated for 12 weeks at 37°C) for non-*M. ulcerans* mycobacterial contamination, and (5) MGITs incubated for up to 12 weeks at 30°C, for rapid demonstration of viability and to re-establish a typical clumping phenotype. DNA extraction (DNeasy PowerSoil Pro Kit, Qiagen) confirmed the presence of *M. ulcerans* by *IS2404* polymerase chain reaction (PCR) as described previously [18]. Endotoxin testing was performed using lyophilized amebocyte lysate (Pierce Chromogenic Endotoxin Quant Kit, Thermo Fisher, Rockford, Illinois, USA).

### Assessing inter-vial variability

To enumerate any differences in CFU count between cryovials prepared in the same batch, *M. ulcerans* JKD8049 was cultured in an orbital shaking incubator for 6 weeks and filtered as described previously. The filtrate was collected in a sterile 30 mL universal container, and then transferred by micropipette into 20% glycerol storage medium (GSM) at a 1:3 ratio. The filtrate/GSM mixture was then vortexed for 5 seconds. The mixture was drawn into a sterile 10 mL stepper pipette, and transferred into 1 mL screw-top cryovials using 300 μL aliquots. The samples were briefly agitated to remove any bubbles and to further mix the sample. Using this process, 46 cryovials (replicating a ‘working cell bank’) of filtered JKD8049 were created from one shaking culture. The cryovials were then cryopreserved at -80°C. After 48 hours, 10% of the vials were thawed on ice for 15 minutes, vortexed for 5 seconds, then spotted onto 7H10/OADC agar in 20 μL volumes. Serial dilutions were made using 20 μL of sample, mixed with 180 uL of PBS in series; 20 μL of the mixtures were spotted. Agar plates were incubated at 30°C for 12 weeks, with CFUs manually counted every two weeks.

### Challenge dose mixing technique

To enumerate bacterial CFU for a prepared challenge dose, a 1 mL low dead-space (LDS) syringe (B-Braun Omnifix-F, Melsungen, Germany), was connected to a 36G drawing-up needle, and 200 uL volume of thawed JKD8049 filtrate was aspirated after vortexing for 5 seconds. 800 uL of PBS was drawn into a separate 1 mL LDS syringe, and the two syringes were connected via luer-lock connector (B-Braun Fluid Dispensing Connector, Melsungen, Germany). The bacterial solution and PBS were then mixed into one another 15 times to create a well-mixed suspension, pressing in a steady back-and-forth motion to compound the suspension (Fig. 2). The solution was then plunged into a single syringe and the luer-lock connector was replaced by a 30G low dead-space hypodermic needle (TSK LDS-30013, Tochigi-Ken, Japan). From this final 1 mL sample, 0.1 mL replicates were injected directly onto 7H10 Middlebrook agar with OADC, creating 10 technical replicates.

**Fig. 2.**
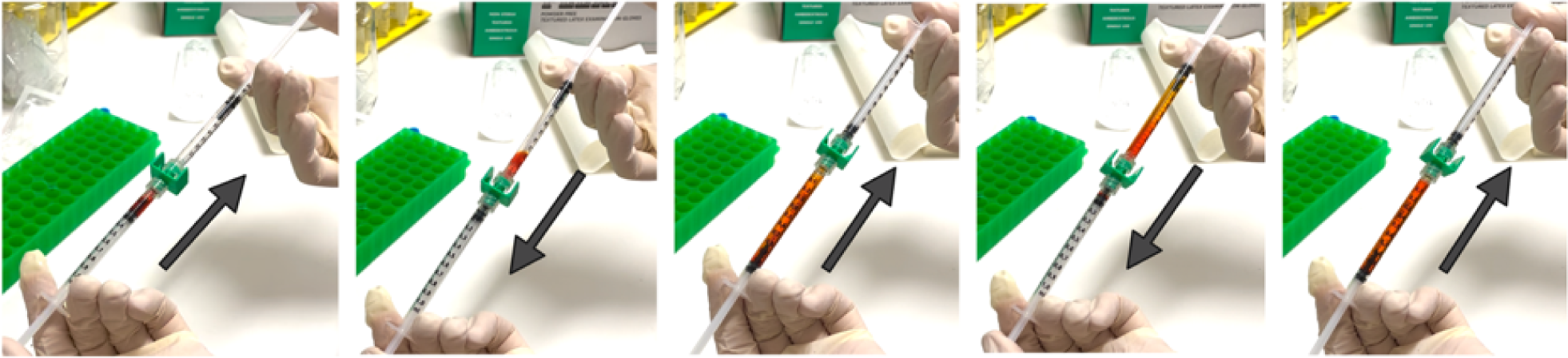
Procedure for diluting cryopreserved cell banks of *M. ulcerans* in phosphate-buffered saline. 200 uL of red dye (bottom syringe, representing a thawed cell bank) and 800 uL of phosphate-buffered saline (top syringe) are connected using a luer lock connector and mixed by repetitive plunging. Four mixing steps are shown. This is performed a total of 15 times to mix samples thoroughly, creating a homogeneous dilution. Arrows represent the direction of plunging to mix the sample seen in the image.

### Mycolactone extraction and detection

Mycobacteria were cultured in SMVT liquid media in stationary flasks for at least 12 weeks. Cultures were then transferred into 50 mL Falcon tubes and spun at 3,824 *g* for 10 minutes at room temperature. The pellet was transferred into a glass tube and weighed. A 10 mL volume of chloroform and methanol (in 2:1 ratio) was added, and the tube was covered with aluminium foil and gently mixed for 5 hours. A 2 mL volume of 0.9% NaCl (w/v) was added and shaken briefly by hand. The sample was then spun at 106 *g* for 10 minutes. The supernatant was removed, and the lower layer was transferred into a fresh glass tube. This extract was dried down under N_2_ on a heat block (set to 45°C) and 2 mL acetone was added. The sample was refrigerated at 4°C overnight, after which the sample was spun at 106 *g* for 10 minutes, and the supernatant transferred to a fresh glass tube. This was again dried under N_2_, leaving acetone soluble lipids, which were analysed using mass spectroscopy. *M. marinum* cultured in SMVT was used as a negative control.

### Macrophage culture and bacterial infection

In order to evaluate the impact of any pre-formed mycolactone that had accumulated in the media during shaking culture, and to understand the immunological implications of de-clumping *M. ulcerans*, human peripheral blood mononuclear cell (PBMC) derived macrophages were exposed to the liquid culture *in vitro*, immediately before and after filtration. Human PBMC derived macrophages were generated as previously described [19]. JKD8049 was incubated in orbital shaking SMVT cultures and filtered (as described earlier). JKD8049 culture samples were collected immediately before and after filtration, and 400 μL of culture material was spiked into 2 mL of macrophage medium with 5 x 10^5^ human macrophages and incubated at 37°C. The multiplicity of infection (MOI) was approximately 1:1, and SMVT media alone was used as a negative control. Stimulated supernatant samples were obtained at serial timepoints during 7 days of incubation and stored at 80°C until analysis by multiplex ELISA (Cytokine & Chemokine 34-Plex Human ProcartaPlex™ Panel 1A) as per manufacturer’s instructions. Cytokine concentrations were calculated using Bio-Plex Manager 5.0 software (Bio-Rad Laboratories, USA) with a five-parameter curve-fitting algorithm applied for standard curve calculations.

### Statistical analysis

Statistical analysis was performed using GraphPad PRISM (version 10.0.1). Continuous variables are reported as mean/standard deviation or 95% confidence interval, as relevant. Comparisons of means between groups were performed using Student’s t test or one-way ANOVA, as appropriate. Results were deemed statistically significant if p < 0.05.

## Results

A collection of five genetically diverse *M. ulcerans* clinical isolates were selected (Table 1) to compare strain-specific antibiotic susceptibility, culture and cryoviability characteristics across all three *M. ulcerans* lineages. JKD8049 was selected for further characterisation in subsequent experiments, in view of its favourable antimicrobial susceptibility pattern. JKD8049 is phylogenomically positioned within lineage 3 of the mycolactone producing mycobacteria complex, and is near identical to other clinical isolate genomes from southeastern Australia (Fig. 3).

**Fig. 3.**
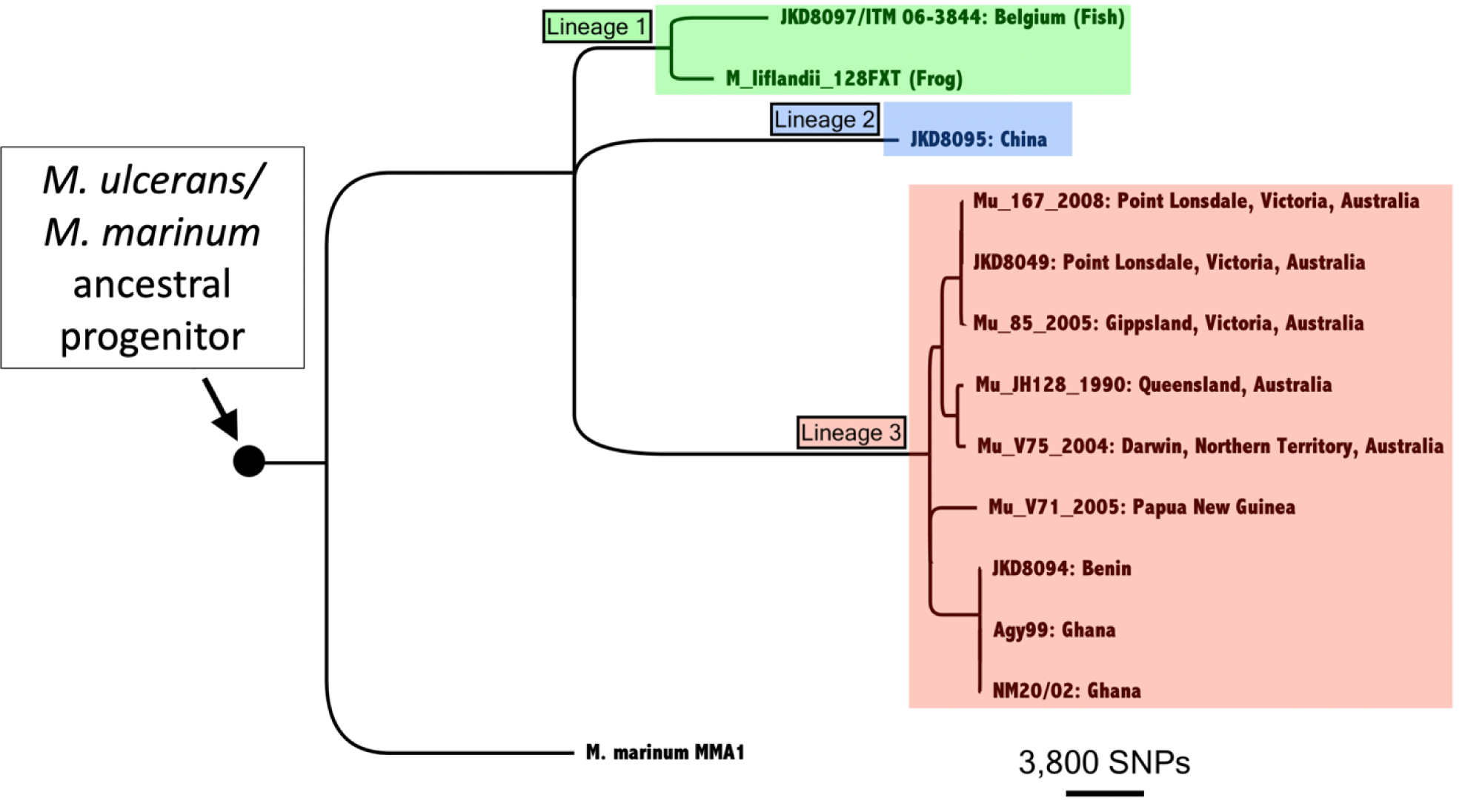
Phylogenomic tree of the *M. ulcerans/M. marinum* complex illustrating the relationships among the isolates examined in this study. The tree, constructed via maximum likelihood based on whole genome SNPs, is rooted with *M. marinum* as an outgroup. The three major lineages within the mycolactone producing mycobacteria complex are highlighted. The horizontal bar provides a scale to represent the number of core SNPs defining branch lengths.

### M. ulcerans JKD8049 is susceptible to all clinically relevant antibiotics

The antibiotic regimen recommended by the World Health Organization for the treatment of BU is rifampicin in combination with clarithromycin [20]; fluoroquinolones (such as ciprofloxacin or moxifloxacin) can be used when clarithromycin is unsuitable [21]. Antibiotic susceptibility testing using a customised and validated MGIT-based method showed strain JKD8049 was susceptible to all three clinically relevant antibiotics tested. Strain 98-0912 (China) and 06-3844 (Belgium) were resistant to rifampicin and ciprofloxacin, and isolate NM20/02 (Ghana) was resistant to ciprofloxacin (Table 2). Based on these results, JKD8049 was the preferred strain for subsequent evaluation.

**Table 2.**
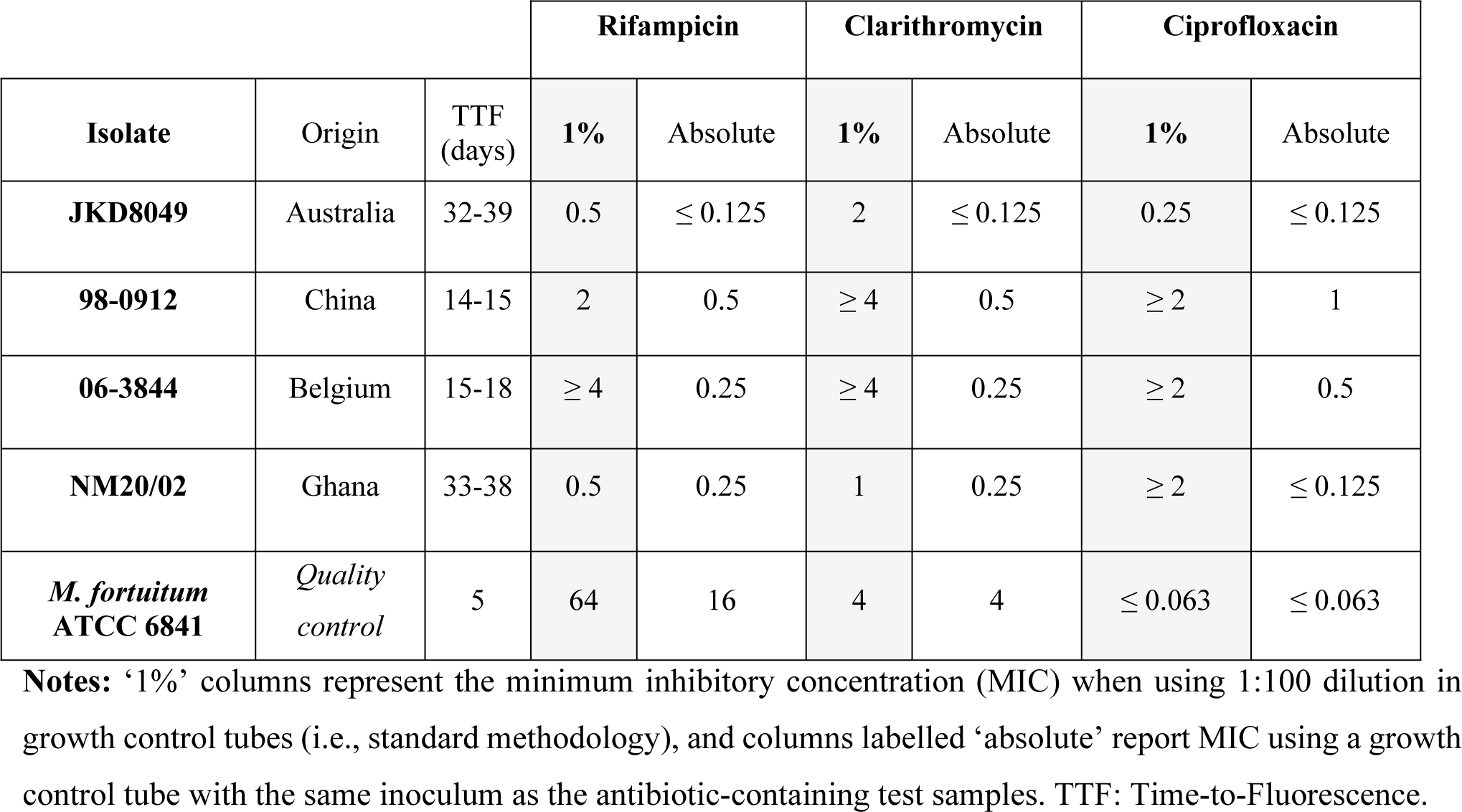
Minimum inhibitory concentrations (μg/mL) of *M. ulcerans* isolates.

| Isolate | Origin | TTF<br>(days) | Rifampicin |  | Clarithromycin |  | Ciprofloxacin |  |
| --- | --- | --- | --- | --- | --- | --- | --- | --- |
|  |  |  | 1% | Absolute | 1% | Absolute | 1% | Absolute |
| JKD8049 | Australia | 32-39 | 0.5 | ≤ 0.125 | 2 | ≤ 0.125 | 0.25 | ≤ 0.125 |
| 98-0912 | China | 14-15 | 2 | 0.5 | ≥ 4 | 0.5 | ≥ 2 | 1 |
| 06-3844 | Belgium | 15-18 | ≥ 4 | 0.25 | ≥ 4 | 0.25 | ≥ 2 | 0.5 |
| NM20/02 | Ghana | 33-38 | 0.5 | 0.25 | 1 | 0.25 | ≥ 2 | ≤ 0.125 |
| *M. fortuitum*<br>ATCC 6841 | Quality<br>control | 5 | 64 | 16 | 4 | 4 | ≤ 0.063 | ≤ 0.063 |

**Notes:** ‘1%’ columns represent the minimum inhibitory concentration (MIC) when using 1:100 dilution in growth control tubes (i.e., standard methodology), and columns labelled ‘absolute’ report MIC using a growth control tube with the same inoculum as the antibiotic-containing test samples. TTF: Time-to-Fluorescence.

### JKD8049 can be cultured in a non-toxic, animal-free medium without genetic or chemical modification

The medium selected for *M. ulcerans* culture was Sauton’s medium (SM) because it is free of animal products (thereby eliminating the risk of transmissible spongiform encephalopathy) and has an established history of use for the culture of the live-attenuated tuberculosis vaccine strain *M. bovis* BCG [22]. Previous studies have also demonstrated that *M. bovis* BCG retains virulence properties when cultured in this medium compared to other media [22], and studies have also demonstrated that *M. ulcerans* retains its ability to produce mycolactone in this medium [23]. Although often routinely added in a research setting to reduce clumping, no surfactants (e.g., polysorbate/Tween) were included; this aimed to minimise chemical modification, particularly considering the presence of hydrophobic lipid-rich structures, including mycolactone, in *M. ulcerans.* As a growth supplement, the vegetable peptone broth ‘Veggietones’ (SMVT) was selected to avoid the use of animal proteins, and because of its established history of use in another bacterial CHIM [24].

All *M. ulcerans* isolates grown in SMVT to stationary phase demonstrated large, clumped growth, with minimal background turbidity, and biofilm formation along the inside of the flask (Fig. 4A). These large clumps, which contain thousands of bacilli (see Fig. S1), would significantly overdose participants if unintendedly injected, as a dose-dependent relationship has been reported in mice [5]. Standardising the dose received by participants will also be important to eliminate differences in groups due to variable dosing. Unfortunately, using 5 μm pore filters to remove these clumps resulted in a low CFU yield in the filtrates of stationary cultures (Figures 4B and 4C).

**Fig. 4:**
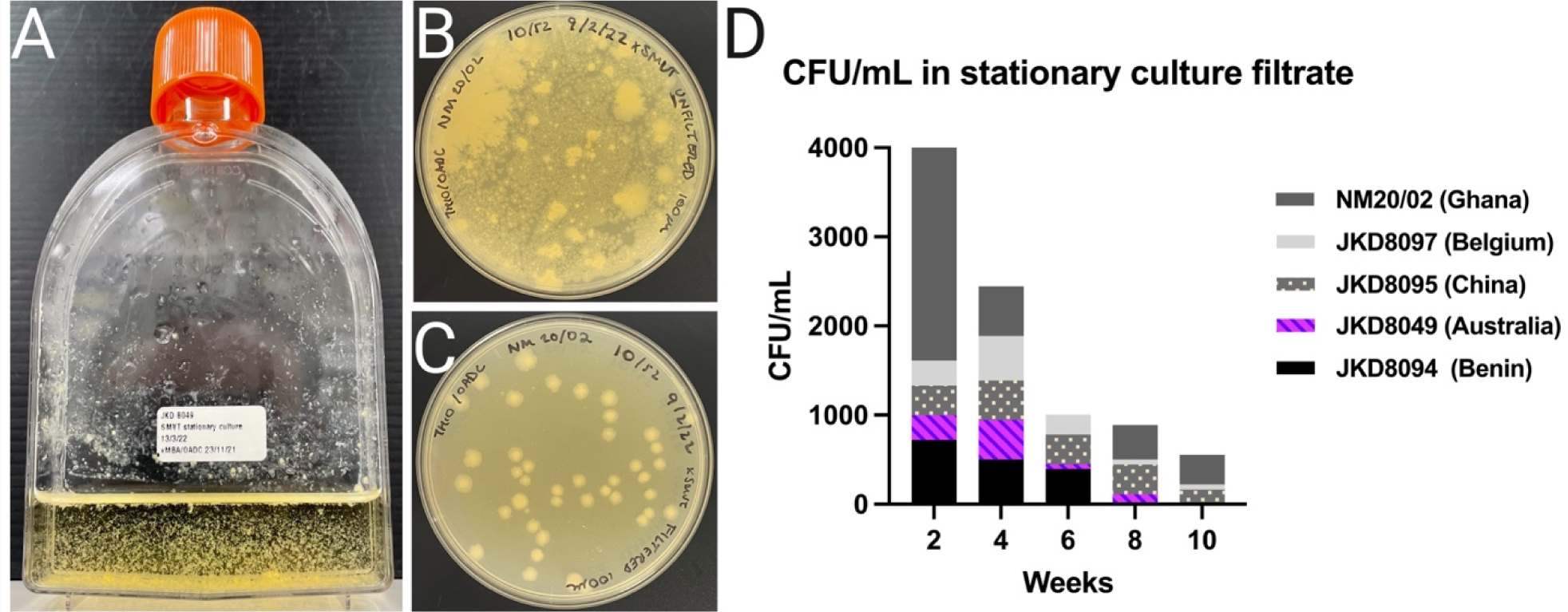
Growth characteristics of *M. ulcerans* in SMVT. (A) JKD8049 cultured for 12 weeks in SMVT, photographed after brief agitation; the growth is in large, dense clumps that rapidly settle with gravity, leaving a background with minimal turbidity, and visible biofilm; (B) 100 μL spread plate of *M. ulcerans* NM20/02 direct from stationary culture in SMVT after 10 weeks of incubation at 30°C; (C) filtrate of NM20/02 shows evenly distributed colonies of approximately similar size and shape in the filtrate; nevertheless, loss of CFU is demonstrated. (D) Enumeration of isolates from stationary cultures in SMVT media and filtered to remove clumps. The upper border of each isolate represents the mean CFU/mL obtained in the filtrate. Enumeration was poor from all isolates except NM20/02 after 2 weeks. Recovery of cells from the filtrate reduced over time, despite visible, clumpy growth in the media.

### Optimising culture conditions for homogeneous growth

Our hypothesis to explain the low yield following filtration after stationary incubation (Fig. 4D), despite the visible and characteristic growth in SMVT, was the presence of innumerable macroscopic clumps, which formed over time and likely saturated the filter pores, thereby obstructing the flow of smaller clumps and/or individual bacilli into the filtrate. To overcome this, orbital shaking cultures (OSC) at 200 rpm were established with 30 mL universal containers and 6 mL SMVT, inoculated with a 10 μL loopful of culture material scraped directly from solid media. In this shaking media, the culture material of some strains was observed to float with minimal disruption due to the hydrophobic nature of the bacteria. This prompted the addition of 20 to 25 sterile 3 mm diameter glass beads to each container with constant agitation to try and create a more homogenous culture suspension and reduce clump size, thereby enabling subsequent efficient filtration. In the containers containing glass beads, the inoculated sample became visibly less clumped within 48 hours of incubation (Fig. 5B), and obvious turbidity became apparent over time, in addition to bacterial pellicle growth and biofilm formation. Additional pilot studies using JKD8049, with a smaller but more defined inoculum of 0.5 McFarland, either did not reliably establish growth in this culture platform, or time-to-turbidity was too slow (results not shown). Subsequent experiments were undertaken to identify the optimal duration of incubation with shaking to maximise the yield for the purposes of subsequent filtration to obtain single-cell preparations. To estimate the optimal time to harvest culture material from SMVT shaking cultures with glass beads, six biological replicates of JKD8049 were established using culture material scraped directly from an agar plate surface using a 10 μL loop. Every two weeks, CFU counts from the centre of the shaking culture were performed, after allowing the cultures to settle with gravity for 15 to 20 minutes. This experiment was designed to replicate the methodology used to harvest liquid culture for the purposes of filtration and cryopreservation, so biofilm and surface pellicle clumps were purposefully avoided.

**Fig. 5.**
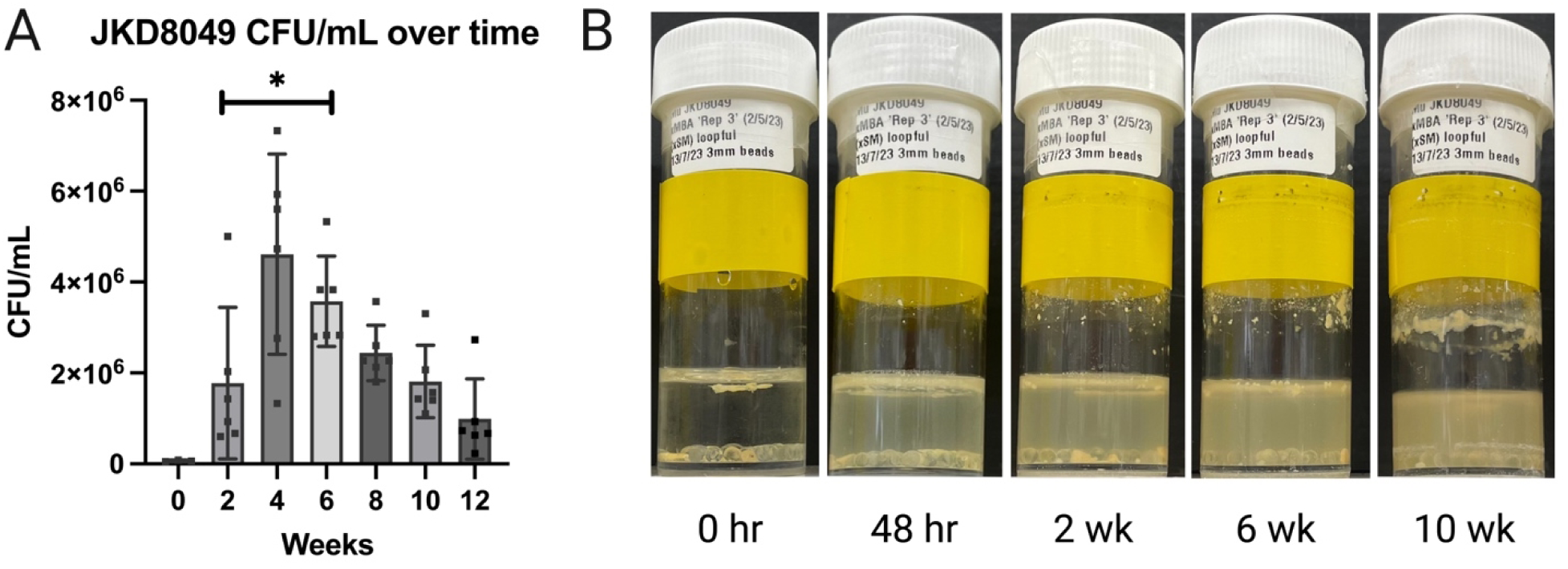
*M. ulcerans* JKD8049 yield in SMVT shaking incubation over time. JKD8049 was cultured in SMVT shaking cultures with 20 to 25 glass beads (3 mm diameter), and samples for CFU enumeration were drawn from the centre of the shaking culture, after pausing shaking for 15 - 20 minutes. (A) represents the average of six biological replicates, except for time 0, which was established from 3 replicates after 48 hours of shaking incubation (Table S1). Mean and standard deviation are shown. Comparison of means was performed by Student’s *t* test, * represents p < 0.05, when comparing 2 weeks’ incubation with subsequent timepoints. Image (B) illustrates visible turbidity, in addition to surface pellicle growth and biofilm formation, in these conditions. The initial inoculated biomass is seen on the far left (note hydrophobic floating material), followed by the disrupted material 48 hours later, with subsequent turbid growth.

As shown in Figure 5A, the yields of well-dispersed JKD8049 in SMVT were greatest when incubated for 4 to 6 weeks. By comparison (Fig. S2), the yield of JKD8049 rapidly decreased when cultured in 7H9/ADC, either due to rapid consumption of nutrients in the medium, or due to biofilm formation. Bacterial yields obtained when JKD8049 was cultured in Sauton’s minimal media were more variable. In summary, harvesting pre-filtration culture material from SMVT shaking cultures was shown to be a reliable method of obtaining a well-suspended sample for subsequent filtration.

### JKD8049 CFU can be enumerated with accuracy to ensure precise challenge dosing

After shaking with glass beads for 12 weeks, CFU counts were performed before and after filtration through a 5 μm diameter filter (Fig. S1), allowing an estimate of the CFUs lost during filtration following this methodology (Table S2). Ziehl-Neilson stains were performed before (Fig. S1C) and after (Fig. S1D) filtration. All filtrate samples demonstrated evenly distributed individual bacilli without any visible clumping. Bacilli were 1.0 to 1.5 μm in length, although longer, dividing bacilli were also occasionally visible. There was some loss of CFU across all isolates de-clumped and enumerated using this methodology (74% loss for JKD8049) (Fig. 6). Nevertheless, filtrates contained an adequate bacterial yield for the purposes of challenge dose preparation from all isolates. We also observed very high accuracy in enumerating CFU in the filtrate of JKD8049, with narrower confidence intervals following filtration than other *M. ulcerans* strains tested (Fig. 6).

**Fig. 6:**
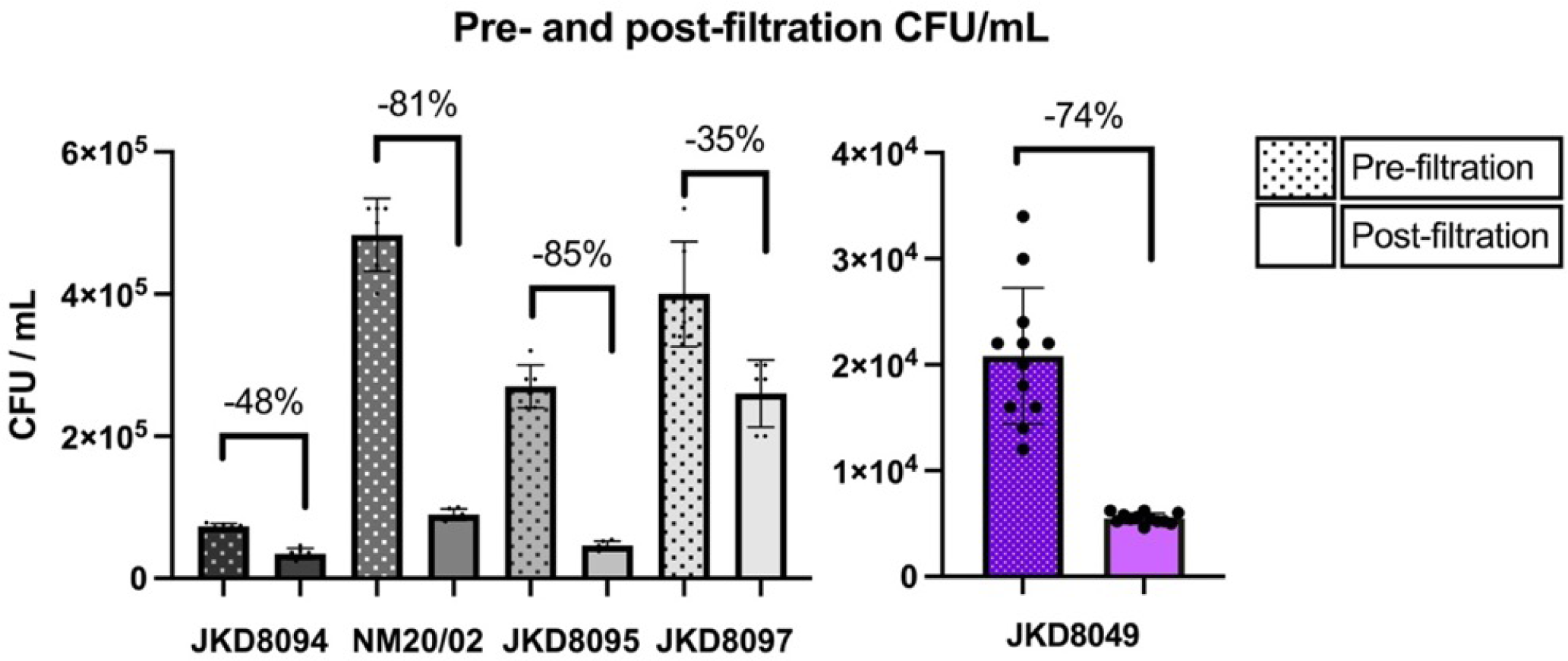
CFU/mL of various *M. ulcerans* isolates after shaking incubation and filtration. The upper border of each isolate represents the mean CFU/mL obtained; error bars represent standard deviation. Pre-filtration (left columns) and post-filtration (right columns) for each strain demonstrate a loss of CFUs from 35 to 85%. On the far right, the closely-spaced replicates (round spots) illustrate the improved accuracy of CFUs obtained after filtration (JKD8049 shown in purple).

### JKD8049 retains viability after cryopreservation

Immediately after filtration, cultures were stored in 20% glycerol (1:3 dilution) and cryopreserved. After 2 to 28 days of cryopreservation, samples of all isolates were placed on ice to thaw slowly until no crystals were visible. Samples were then vortexed for 5 seconds and enumerated. To determine the viability of samples over time after thawing, the experiment was performed at three timepoints: immediately after thawing, then after 1 hour and 2 hours resting on ice. A longer viability timeframe was evaluated for JKD8049, with viability tested after 2 and 4 hours on ice. This demonstrated that isolates NM20/02 and JKD8049 retained viability following cryopreservation (Fig. 7, Table S3).

**Fig. 7:**
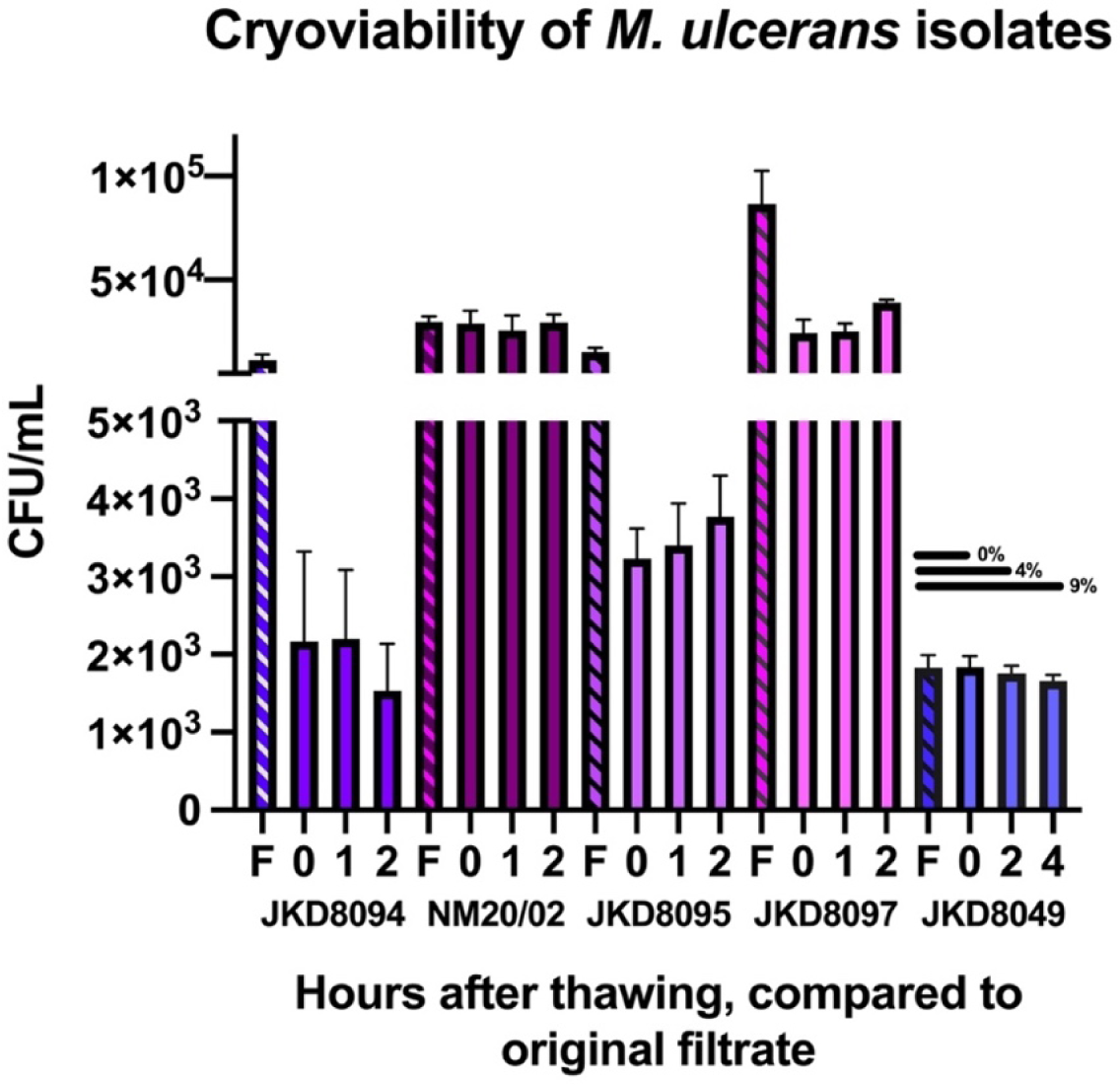
CFU/mL of various *M. ulcerans* isolates after filtration and cryopreservation in glycerol at -80°C (accounting for dilution factor) compared to CFU/ml expected from original filtrate (F) sample. CFU/mL were counted immediately after thawing (0 hours), and at two additional timepoints. The upper border represents the mean CFU/mL obtained; error bars represent standard deviation. Viability loss (%), is shown for JKD8049, others are available in Table S3.

To estimate the potential difference between the number of *M. ulcerans* bacilli in the cryopreserved vial and the observed final CFU count, 5 μL volumes were drawn from the thawed sample of isolate JKD8049, in triplicate, and a Ziehl-Neilson stain was performed after heat-fixing to a glass slide. In each 5 μL volume, the whole spot was manually scanned at high power, and 58, 58 and 50 bacilli were counted (mean 55.3 bacilli per 5 μL volume). When compared to the actual CFU count observed in 5 μL (9.2 CFU), this suggests that approximately 83% of the bacilli seen in the sample were non-viable. Given that there was no difference in the viable cell count between the cryopreserved sample and the filtrate CFU count (accounting for the dilution factor), it is likely that the visualised cells were non-viable before cryopreservation and had likely become non-viable during orbital shaking. This may further support using a duration of 4 to 6 weeks of shaking incubation rather 12 weeks, to minimise bacterial death due to nutrient exhaustion during shaking culture.

### JKD8049 cryopreservation viability does not require additional supplementation

Given that the local standard cryopreservative contains tryptone soya broth, samples of JKD8049 were tested using ‘Veggietones’ replacing the tryptone soya broth (at the same concentration), compared to glycerol without any supplemental agent. This demonstrated that, at all timepoints, there were significantly higher CFU in samples cryopreserved without supplemental Veggietones. There was no statistically significant difference (p >0.05) between any of the timepoints within the ‘supplement’ and ‘no supplement’ groups (Fig. 8A).

**Fig. 8.**
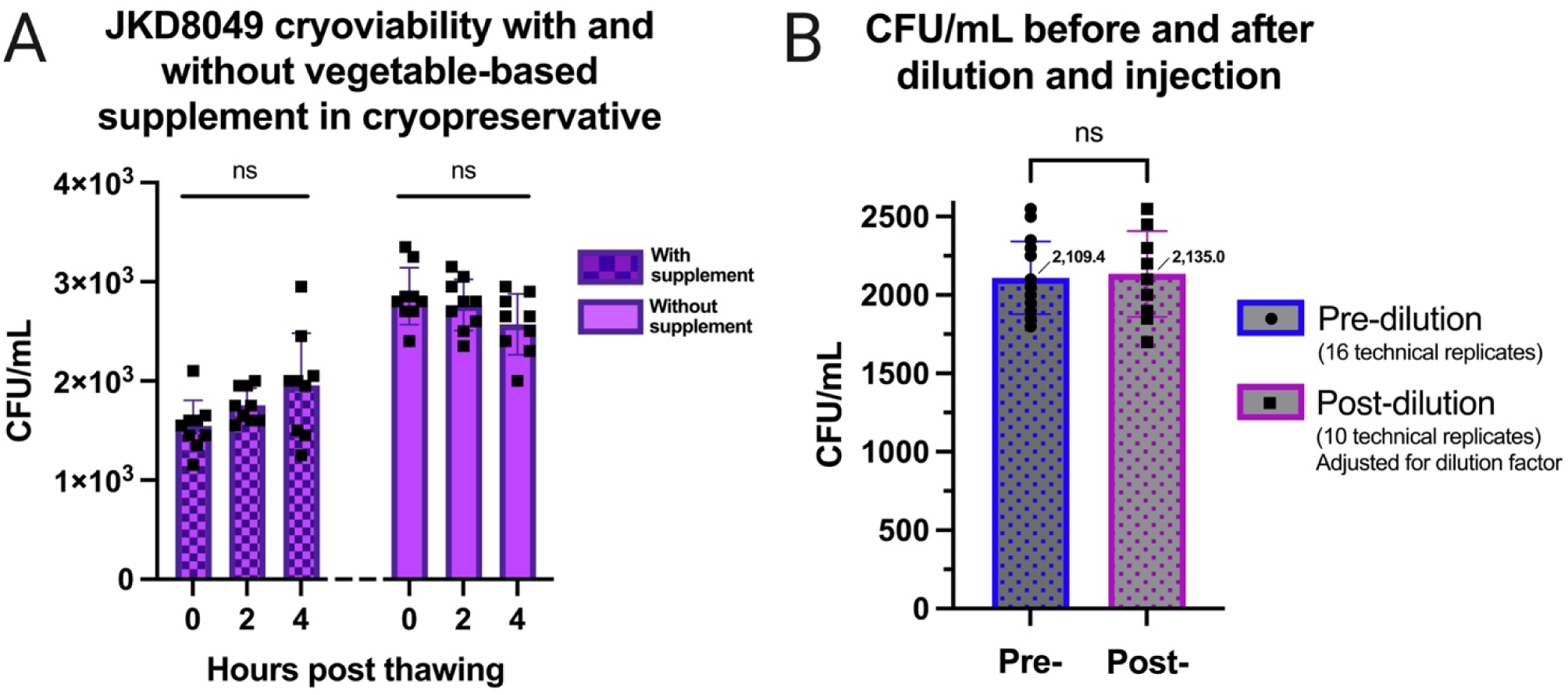
(A) JKD8049 cryopreserved with and without supplement. Filtered JKD8049 was stored in either glycerol with ‘Veggietones’ supplement (left columns) or glycerol without supplement (right columns). Samples were tested immediately after the sample was thawed on ice (‘0 hours’) or at subsequent timepoints (‘2 hours’ and ‘4 hours’). The upper border of each isolate represents the mean CFU/mL obtained; error bars represent standard deviation. **(B) CFU/mL obtained before and after dilution using needle and syringe system**. There was no significant (ns) difference in CFU before (left) or after (right) dilution in PBS and injection through 30G needle and LDS syringe (mean and standard deviation are shown, comparison of means was performed using Student’s t test).

### JKD8049 cell banks can be created with minimal inter-vial variability

The process described above was used to generate 46 single-use cryopreserved vials containing *M. ulcerans* JKD8049. To ensure a consistent and predictable dose of *M. ulcerans* is obtained when selecting from such a bank of prepared cryovials, 10% of the samples were thawed, taking every 10^th^ vial to ensure a representative sample across the manufactured lot. Samples were vortexed briefly and spotted in serial dilution for CFU enumeration. This demonstrated no significant difference in the CFU counts obtained between the cryovials (one-way ANOVA, p=0.13) (Fig. 9).

**Figure 9:**
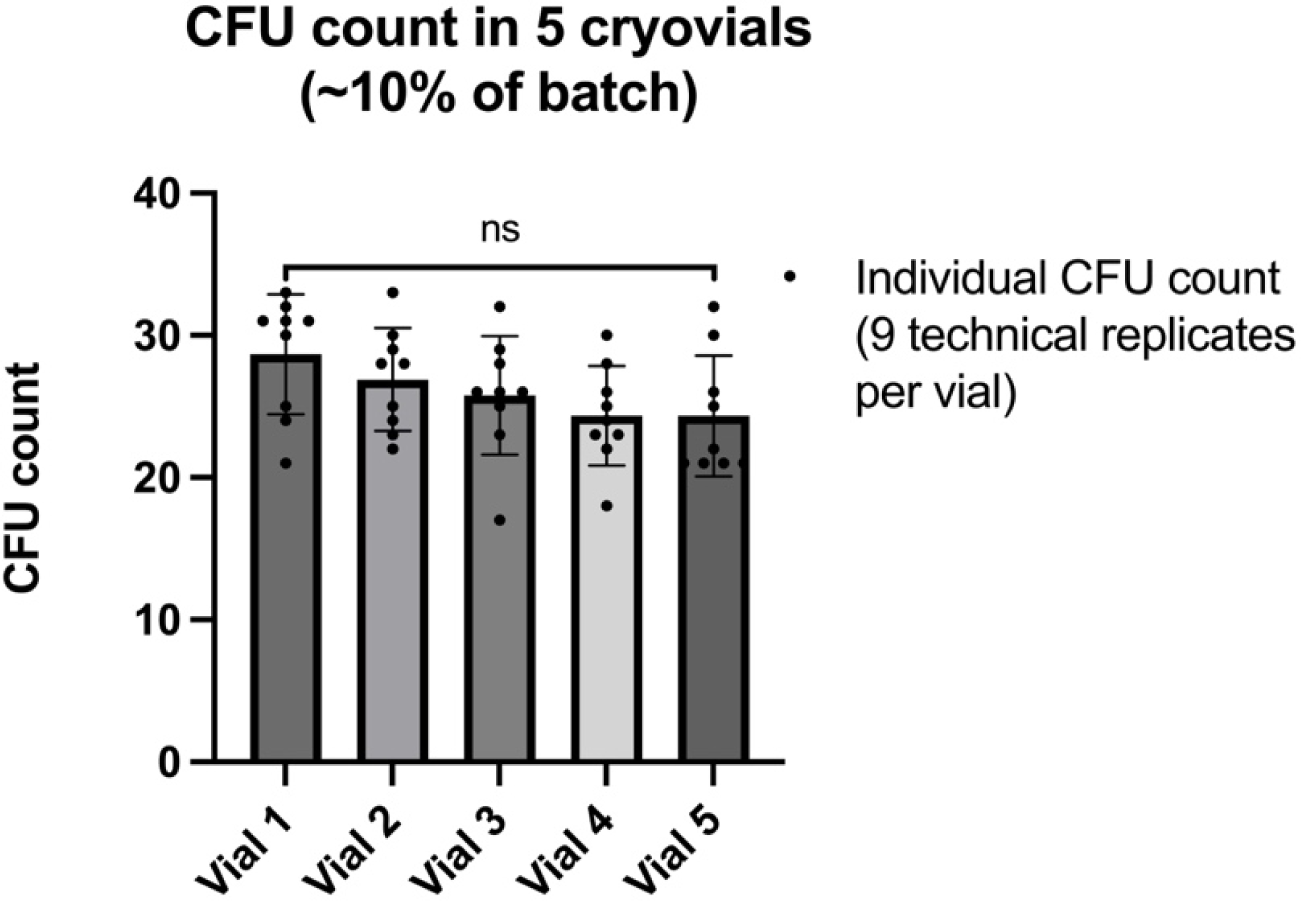
CFU counts obtained from five vials of cryopreserved *M. ulcerans* JKD8049 from a cell bank of 46 cryovials. There was no significant difference in CFU count across all five vials tested (presented here as CFU count in each technical replicate); mean and standard deviation are shown (one-way ANOVA, p = 0.13).

### Filtered, cryopreserved JKD8049 can be diluted to low doses for parenteral injection

To confirm that the CFU count is conserved after (i) thawing, (ii) diluting to the required low dose and (iii) injecting using needle and syringe, CFU counts were performed before and after sample preparation, replicating the methodology anticipated to be implemented for challenge administration. Initial CFU count was performed using a thawed preparation of cryopreserved, filtered JKD8049, which had been cultured for 12 weeks, as described earlier. Sixteen technical replicates, each using 20 μL spot volumes, were used to determine the CFU count before sample preparation. Post-dilution samples were spotted in 0.1 mL volumes, through a low dead space needle and syringe directly onto 7H10/OADC agar and incubated for 12 weeks. 0.1 mL volume was selected, as it replicates the dose anticipated for human challenge. Accounting for the dilution factor, there was no difference in the mean CFU count measured, suggesting no significant CFU loss using this technique (Fig. 8B).

### Quality control

Using a range of selective and non-selective media, 10% of cryovials produced in a single cell bank were tested for adventitious agents, with no bacterial or fungal contamination identified. Endotoxin testing prior to dilution demonstrated an endotoxin level (0.118 EU/mL) far below the regulatory endotoxin limit for this product (K/M = 1,750 EU/mL, based on a cell bank volume of 0.2 mL and an average 70 kg person) [25], supporting the pyrogen-free nature of the challenge product. Culture of de-clumped bacilli in 7H9/OADC MGIT media demonstrated characteristic clumpy growth in all strains tested, with acid-fast bacilli confirmed on Ziehl-Neelsen staining.

### JKD8049 produces mycolactone after in vitro culture in SMVT

To study the true immunobiology of Buruli ulcer in a generalisable model of infection, the selected isolate must still produce the main virulence factor, mycolactone. It is crucial that mycolactone is further characterised because minor sequence modifications in the ML genes confer a variety of structural variants. ML A/B, produced by African strains, is the most cytotoxic, while the potency is attenuated in ML C (produced by Australian strains) and the other structural variants [26,27]. Australian *M. ulcerans* strains also produce a fraction of ML A/B, which may be more important for the pathogenesis than ML C in Australian isolates [27]. *M. ulcerans* is reportedly able to produce ML in Sauton’s media [23]. Compared to 7H9 Middlebrook broth, Sauton’s media has been shown to induce *M. bovis* BCG to acquire properties associated with virulence, with increased ability to withstand intracellular conditions and modulate immune responses [22]. Therefore, lipids were extracted from *M. ulcerans* cultured in SMVT and used to determine the presence of mycolactone using liquid chromatography–mass spectrometry. All isolates in this study were confirmed to produce mycolactone when cultured in SMVT, including JKD8049, which produced mycolactone C and A/B, as anticipated for Australian isolates.

### Evaluating the role of metabolic shift on bacterial growth

Considering the importance of precisely enumerating CFU count to prevent under- or overdosing participants, a series of experiments were designed to interrogate the conditions which may improve the growth of *M. ulcerans* JKD8049 colonies on solid media and therefore improve CFU yield.

To investigate the possibility that CFU count may be improved by minimising the metabolic shift in nutritional conditions when plating onto solid media, JKD8049 was cultured in SMVT liquid media in shaking cultures for 12 weeks, then spotted onto SMVT converted into a solid media with agar, alongside 7H10/OADC agar as a control. Enumeration was performed using 5 μm spots, with six technical replicates per plate, in serial dilution. SMVT was used as the diluent for all serial dilutions on both agar plates. After 12 weeks, plates were photographed and are presented in Figure 10. When spot plated on SMVT agar, JKD8049 demonstrated luxurious growth, with the neat (undiluted) sample producing large, spreading colonies, particularly along the walls of the square plate (Fig. 10B). However, there was no growth at any dilution (i.e., lower inoculum). This may explain earlier observations that a low inoculum of JKD8049 could not reliably establish a turbid culture in SMVT, but a heavy inoculum (i.e., 10 μL loopful of scraped culture material, ≥ 1x10^9^ CFU) reliably established a turbid culture in this media. Additionally, this phenomenon of spreading growth appeared to be facilitated by the supplementation of Sauton’s media with ‘Veggietones’, allowing the rapid growth of individual colonies (Fig. 10C), a phenomenon not apparent on Sauton’s minimal media (Fig 10D; seen after 12 weeks of incubation at 30°C).

**Fig. 10:**
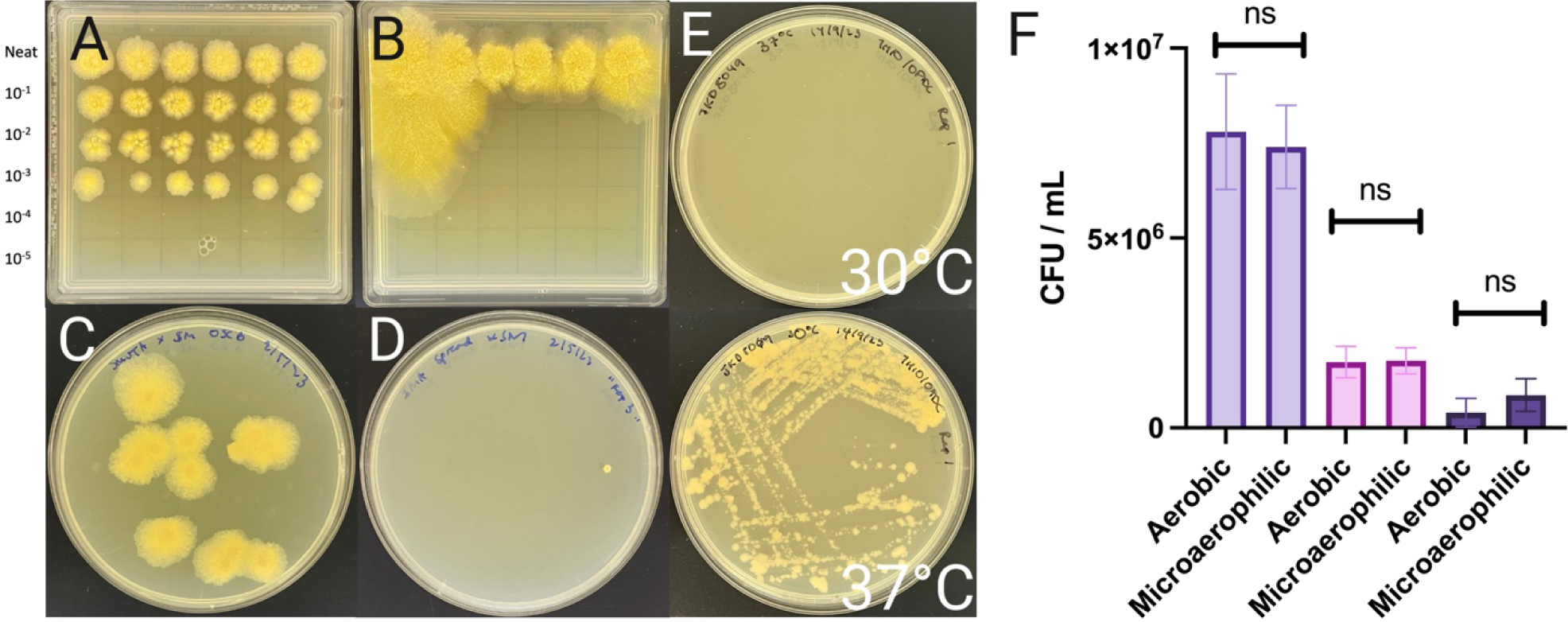
Comparisons of JKD8049 in various culture conditions. Image (A) shows 5 μL spot plates of JKD8049 on 7H10/OADC agar compared to (B) SMVT agar. Image (C) shows 100 μL spread plates of *M. ulcerans* JKD8049 spread onto SMVT agar, compared to (D) Sauton’s minimal media agar. Although triplicates were produced of each agar, only one colony was visible from all three Sauton’s media agar plates, whereas colonies on all three SMVT agar replicates demonstrated spreading growth. Images A-D were captured after 12 weeks of incubation at 30°C. (E) *M. ulcerans* JKD8049 cultured at 30°C (top) and 37°C (bottom) confirmed thermal restriction of this isolate. (F) Shown are mean and standard deviation of biological triplicate experiments of *M. ulcerans* JKD8049 cultured in both aerobic and microaerophilic conditions.

### JKD8049 is unable to grow at core human body temperature

Subacute haematogenous spread of *M. ulcerans* has been occasionally reported primarily in African populations, where it is speculated that isolates are more thermotolerant than those from temperate regions [26]. To assess thermotolerance, *M. ulcerans* JKD8049 was streaked onto 7H10/OADC agar in triplicate, and plates were cultured at 37.0°C (SD: 0.15°C) and 30.0°C (SD: 0.48°C), as monitored using a portable electronic thermometer. Plates were reviewed after 12 weeks to evaluate growth (Fig. 10E), confirming that isolate JKD8049 is unable to grow at 37°C [28].

### JKD8049 does not require a microaerophilic environment for growth

To investigate the possibility that CFU count may be improved by culturing in microaerophilic conditions, as suggested by previous studies [29], three biological replicates of *M. ulcerans* JKD8049 were cultured in SMVT using shaking incubation in aerobic conditions. After 4 weeks, samples were spotted onto 7H10/OADC agar and incubated in either microaerophilic or aerobic conditions. Plates were examined once after 6 weeks of incubation; fortnightly review of CFU was purposefully avoided to minimise the impact of introduced oxygenation. This demonstrated that there was no difference in mean growth between conditions (p < 0.05, Student’s *t* test) (Fig. 10F).

### JKD8049 has a conserved repertoire of genes encoding candidate vaccine antigens

To ensure that the isolate is fit-for-purpose, it must be capable of expressing a range of candidate vaccination antigens. A systematic review of all putative vaccine targets [30] informed the antigens listed in Table 3, and a recent *in silico* study [31] also suggested the importance of including the MFS transporter proteins in the challenge strain. Whole genome sequencing using Illumina technology was used to confirm the presence of a range of candidate vaccine targets (Table 3).

**Table 3.** Candidate vaccine targets and encoding gene presence in JKD8049.

| Candidate target antigen | Code | Gene in JKD8049 |
| --- | --- | --- |
| Ag85A | MUL_4987 | Present |
| Ag85B | MUL_2970 | Present |
| Hsp18 | MUL_2232 | Present |
| Hsp65 | MUL_1393 | Present |
| 21 kDa protein | MUL_3720 | Present |
| Ag85B-EsxH | MUL_1210 | Present |
| Polyketide synthase (*pks*) modules (1-9) | N/A | Present |

### JKD8049 remains genetically stable during manufacture and challenge

To assess the genomic stability of JKD8049 after serial passage, isolate JKD8049 was cultured in successive solid/liquid animal-free media (using SMVT in liquid or in agar), with successive passages over 22 months, to replicate the conditions of challenge dose manufacture. After five passages, five separate populations were selected for whole genome sequencing. From these isolates, sequencing did not identify any SNPs compared to the JKD8049 reference chromosome (GenBank accession: CP085200.1), underscoring a level of genetic stability for this isolate.

### De-clumped JKD8049 stimulates human macrophages ex vivo

To evaluate the ability of JKD8049 to elicit an immune response, and to understand the immunological implications of de-clumping *M. ulcerans*, we exposed human peripheral blood-derived macrophages *ex vivo* with the liquid culture, before and after filtration. Figure 11 illustrates that exposed macrophages induced the secretion of the cytokines IL1b, Il-6, Il-8, and TNFα. Similar responses between treatment of macrophages with filtered and non-filtered culture indicate that pre-formed mycolactone was unlikely to be present in any significant concentration. These results also demonstrated that the media is immunologically inert, similar to the unstimulated control.

**Fig. 11.**
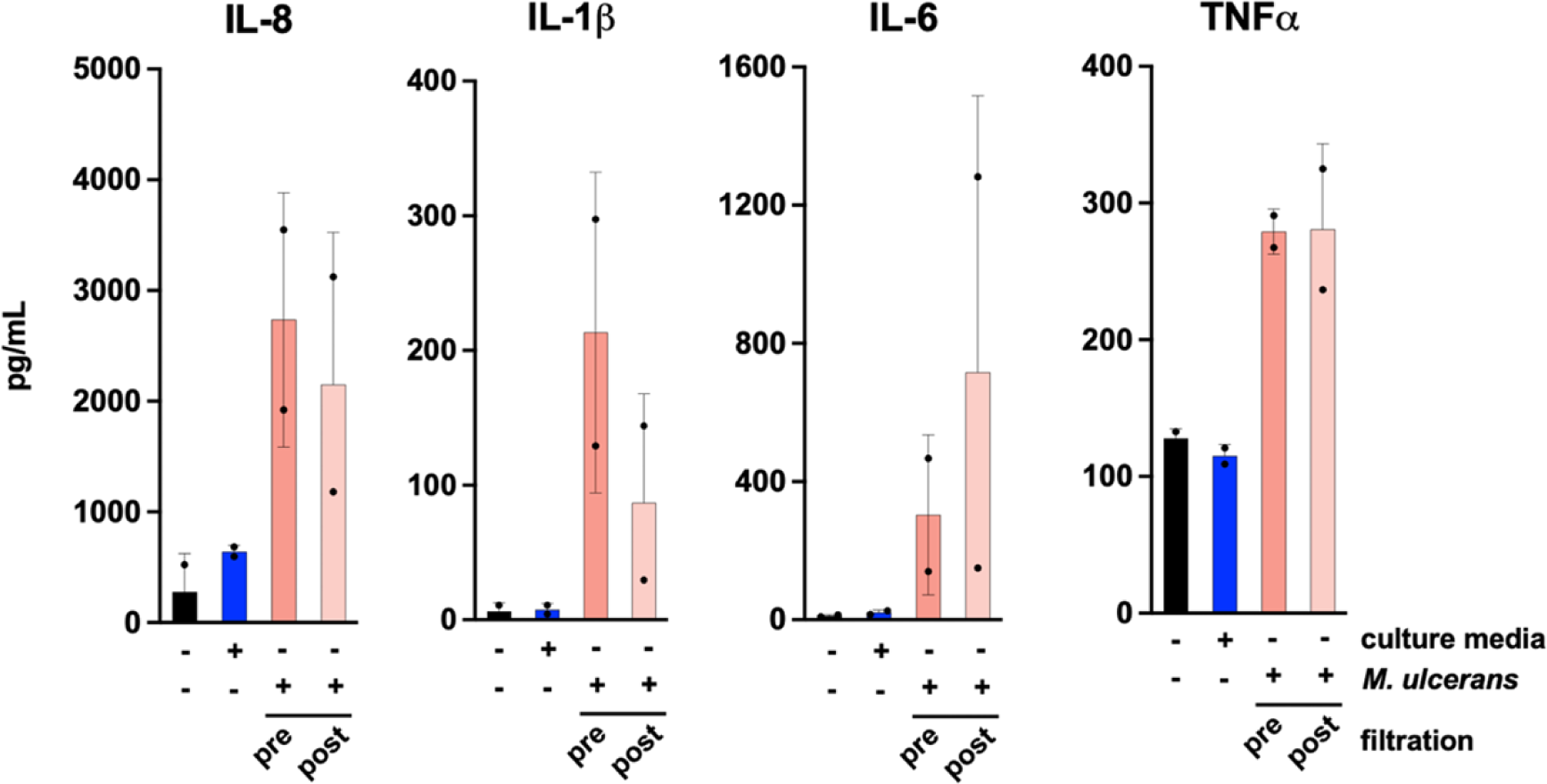
ELISA measurement of IL-8, IL-1b, IL-6 and TNFα secreted by human macrophage in response to 0.4 mL of *M. ulcerans* JKD8049 culture sample, before and after filtration. SMVT media only was analysed in addition to unstimulated controls.

## Discussion

This study has systematically characterised *M. ulcerans* JKD8049 as a candidate isolate for the first controlled human infection model of Buruli ulcer. Notwithstanding studies investigating *M. bovis* BCG vaccine as a proxy for infection with *M. tuberculosis* [32], this work advances what is anticipated to be the first pathogenic mycobacterial human challenge model. This work also demonstrates that the methodology can be implemented across a variety of isolates, if other investigators wish to reproduce this approach for their own challenge agent manufacture, or for any other relevant application.

This study provides an approach to testing *M. ulcerans* antibiotic MIC, which infers susceptibility based on clinical breakpoints, defined by associations such as the Clinical and Laboratory Standards Institute [33]. MIC testing is limited by the lack of standardised methodology, with guidelines suggesting that each laboratory should establish and validate their own methodology [33]. Established CLSI susceptibility breakpoints for slowly growing non-tuberculous *Mycobacteria* (other than *M. avium* complex and *M. kansasii*) suggest that rifampicin MIC ≥ 2 *μ*g/mL should be categorised as ‘resistant’ [33]. Of the clinical isolates tested, strain 06-3844 demonstrated the highest rifampicin MIC; this isolate was first identified as a mycolactone-producing member of the *M. marinum* complex, which typically demonstrate rifampicin MICs within the range of 0.124 – 4.0 *μ*g/mL [34]. This isolate is therefore inappropriate to progress to human use. In their original description, Faber and colleagues reported that the isolate 98-0912 (isolated in China), was rifampicin resistant, which prompted a switch in antibiotic therapy, although they do not describe their methodology or by which criteria resistance was defined [8]. Nevertheless, this strain did demonstrate a relatively high rifampicin MIC compared to the Australian and African isolates, and is indeed considered resistant to rifampicin by CLSI criteria [33]. Strain NM20/02, an attenuated strain originally isolated in Ghana [5], demonstrated rifampicin MIC consistent with previously reported Ghanaian isolates [35]. The Australian isolate JKD8049 demonstrated a rifampicin MIC which was higher than previously reported by Omansen and colleagues, although the methodology used in their study did not use a growth control to establish the MIC [6]. Indeed, using an arbitrary timepoint of 21 days [6] or 28 days [36] of incubation, rather than defining the MIC by comparison to an antibiotic-free growth control, may explain previous discrepancies. Nevertheless, the rifampicin MIC for JKD8049 reported here is categorised as ‘susceptible’ according to CLSI breakpoints [33], is similar to that reported using the agar dilution method [37], and is lower than other Victorian historical reference strains [38].

Established susceptibility breakpoints for *M. ulcerans* suggest that clarithromycin MIC ≤ 8 *μ*g/mL should be categorised as ‘susceptible’ using CLSI criteria (and is representative for newer macrolides, such as azithromycin) [33]. As discussed by Portaels *et al*. [39], a range of clarithromycin MICs were found to be below the peak plasma concentration of clarithromycin obtainable in humans, noting that the administration of 500 mg of clarithromycin twice daily for up to 3.5 days to volunteers results in mean peak plasma concentrations between 2.4 and 3.5 μg/mL [40]. Nevertheless, because clarithromycin has a large volume of distribution [41], it achieves higher concentrations in tissues than in the blood [42]. Additionally, it has excellent intracellular penetration, which is likely to be most useful in early stages of infection (prior to the formation of extracellular clumps) and in intracellular bacteria as they advance centrifugally to propagate infection [43,44]. In their study, Portaels and colleagues tested a broad geographic representation of isolates, and although Australian and Malaysian strains exhibited the highest clarithromycin MICs (between 0.5 to 2 μg/mL), no strain demonstrated an MIC greater than 2 μg/mL [39]. In the present study, isolates NM20/02 and JKD8049 demonstrated clarithromycin MIC values which are both below predicted peak plasma concentrations [40]. It is worthwhile noting that the *in vitro* susceptibility of *M. ulcerans* to clarithromycin is dependent on the pH of the media; in Australian isolates, the clarithromycin MIC ranges from 0.5 to 4 μg/mL at pH 6.6 and < 0.125 – 0.5 μg/mL at pH 7.4 [39]. This study therefore overestimates clarithromycin MIC values, because the pH in 7H9 broth used in MGITs is 5.9 [11].

Established susceptibility breakpoints for *M. ulcerans* suggest that ciprofloxacin MIC ≤ 1 μg/mL should be categorised as ‘susceptible’ using CLSI criteria [33]. Although a fluoroquinolone is unlikely to be required in the treatment regimen in the majority of CHIM participants, this study showed that *M. ulcerans* JKD8049 demonstrated the lowest ciprofloxacin MIC, and was the only tested isolate susceptible by CLSI criteria [33]. In their study, Owusu *et al.* reported that Ghanaian strains demonstrated a mean ciprofloxacin MIC of 1.15 *μ*g/mL [35]. NM20/02, with a ciprofloxacin MIC of ≥ 2 *μ*g/mL, is considered to have at least ‘intermediate’ susceptibility by current criteria [33].

To assess this methodology in the context of discussion surrounding the use of the proportion method [45], growth control tubes without further dilution were also included (i.e., the growth control tube contained the same bacterial inoculum as the antibiotic-containing tube). When comparing MIC values to those previously reported in the literature, the methodology presented here demonstrates the utility in diluting the growth control tube by 1:100 for *M. ulcerans* isolates, enabling improved resolution of MIC values. Using this methodology, JKD8049 has rifampicin and clarithromycin MICs which are similar to those determined with an agar proportion methodology [37]. The proportion method identifies a more conservative (i.e., higher) estimate for the MIC. The MIC values of the quality control (QC) isolate further support quantifying *M. ulcerans* MICs using this platform, although a slow-growing mycobacterial QC isolate will be prioritised in future validation.

A limitation of the lengthy timeframes required for growth is the possibility of drug degradation in the 7H9 broth. When incubated at 37°C, rifampicin has previously been shown to degrade over time, until reaching undetectability at 6 weeks’ incubation, although residual rifampicin (or degradation products) likely continue to inhibit growth [46]. Antibiotic degradation would theoretically result in higher apparent MIC values for slower growing strains; nevertheless, the two slowest growing strains still demonstrated the lowest rifampicin MIC values. These results support the use of the MGIT system for *M. ulcerans* susceptibility testing according to the established framework for *M. tuberculosis* susceptibility testing [11]. The benefits of this system include (1) the ability to automate results, (2) the ease-of-use due to removing the need for CFU counting (and avoiding counting errors), and (3) the shortened turn-around time. The MGIT platform is also a closed system, so after inoculation, contamination is less likely than traditional agar-based susceptibility testing. Nevertheless, the platform is not inexpensive, and requires further validation.

One of the major limitations working with *M. ulcerans* is the bacterium’s slow growth, with a doubling time of ∼48 hours *in vitro* [47]. This leads to long incubation periods, generally up to 12 weeks, within an optimal temperature range of 30 – 33°C. The long incubation period increases the opportunity for microbial contamination, emphasising the importance of performing all experimental work within strictly sterile conditions. Although a more rapid time to culture positivity is an attractive characteristic, it is not necessarily a requirement for a candidate CHIM strain. Sauton’s media with supplemental pea flour-based peptone appears to be a satisfactory animal-free medium to establish turbid growth of *M. ulcerans* JKD8049, using continuous mechanical agitation to minimise clumping without detergent/Tween. Although mechanical de-clumping procedures are often used to create suspensions for nephelometry or other applications (e.g., vortexing with or without glass beads, needle-syringing, and/or settling with gravity), they have variable success in de-clumping *M. ulcerans* with accuracy. Filtration of turbid liquid culture through 5 μm pore filters successfully removes clumps and optimises enumeration accuracy. This filtrate can then be cryopreserved in 20% glycerol with minimal reduction in viability. Completely de-clumping mycobacteria to individual cells appears to minimise sticking of cells to the syringe/needle material, enabling dilution and injection of cryopreserved material. Reassuringly, there was no significant difference in the extent of immunomodulation conferred by mechanically removing *M. ulcerans* clumps. Finally, this report also describes an approach to ensure the purity, potency and identity of the challenge agent.

The observation that *M. ulcerans* JKD8049 grows as a luxurious biofilm on SMVT agar when incubated in a larger inoculum, but not in diluted samples, suggests an unrecognised quorum sensing phenomenon, and warrants further research. Nevertheless, the addition of a vegetable-based supplement appears to improve the growth characteristics of *M. ulcerans* JKD8049, shortening incubation times and therefore reducing the risk of environmental contamination. Conversely, supplementing glycerol storage media with this product seemed to reduce cryoviability, and is therefore not required to improve revival following cryopreservation.

Finally, the absence of SNP variants across multiple passages spanning 22 months suggests that isolate JKD8049 will maintain genomic stability throughout the challenge dose production process. The observed genetic stability, marked by a lack of SNP accumulation, aligns with previous findings from a population genomic study of clinical isolates in southeastern Australia, reporting a slow molecular clock, with a mutation accumulation rate of 0.39 SNPs per chromosome annually (excluding insertion sequence elements) [48]. Given this marked genomic stability, isolate JKD8049 emerges as an exemplary candidate challenge strain.

## Conclusion

In finding that JKD8049 meets the requirements of a BU human challenge strain, we have reported here on strategies to refine determination of *M. ulcerans* antibiotic susceptibility, to optimise turbid culture in an animal-free liquid media, and to systematically de-clump *M. ulcerans* without detergent for accurate, ultra-low dosing. Our findings will inform the manufacturing procedures to establish cell-lot banks for human challenge, in accordance with regulatory standards.

## Acknowledgements

We thank Jean Y.H. Lee, Belinda B. Lin, Alexandra Krause, Sher Maine Tan and Rachel Simmonds for technical advice or support in the laboratory; Gerd Pluschke for provision of isolate NM20/02.

## Supplementary data

**Table S1.** *M. ulcerans* JKD8049 mean CFU/mL in SMVT over time using orbital shaking cultures with glass beads, incubated at 30°C. 6 biological replicates were tested (Rep 1-6).

|  | 48 hours | 2 weeks | 4 weeks | 6 weeks | 8 weeks | 10 weeks | 12 weeks |
| --- | --- | --- | --- | --- | --- | --- | --- |
| Rep 1 | 6.30E+04 | 1.03E+06 | 1.33E+06 | 2.80E+06 | 2.27E+06 | 1.40E+06 | 9.33E+05 |
| Rep 2 | 8.33E+04 | 6.67E+05 | 5.93E+06 | 3.83E+06 | 2.27E+06 | 2.07E+06 | 2.33E+05 |
| Rep 3 | 5.00E+04 | 1.43E+06 | 7.33E+06 | 5.33E+06 | 3.57E+06 | 3.30E+06 | 2.73E+06 |
| Rep 4 | - | 6.00E+05 | 2.76E+06 | 2.83E+06 | 2.60E+06 | 1.43E+06 | 7.33E+05 |
| Rep 5 | - | 9.33E+05 | 4.73E+06 | 3.83E+06 | 1.80E+06 | 1.10E+06 | 6.33E+05 |
| Rep 6 | - | 2.03E+06 | 5.60E+06 | 2.00E+06 | 2.13E+06 | 1.57E+06 | 6.67E+05 |
| Mean | 6.54E+04 | 1.12E+06 | 4.61E+06 | 3.75E+06 | 2.44E+06 | 1.81E+06 | 9.88E+05 |
| SD | 1.68E+04 | 6.89E+05 | 2.33E+06 | 2.01E+06 | 8.93E+05 | 9.79E+05 | 8.48E+05 |
| *p* | N/A | *Comparator* | 0.031 | 0.046 | 0.382 | 0.964 | 0.330 |

**Table S2.**
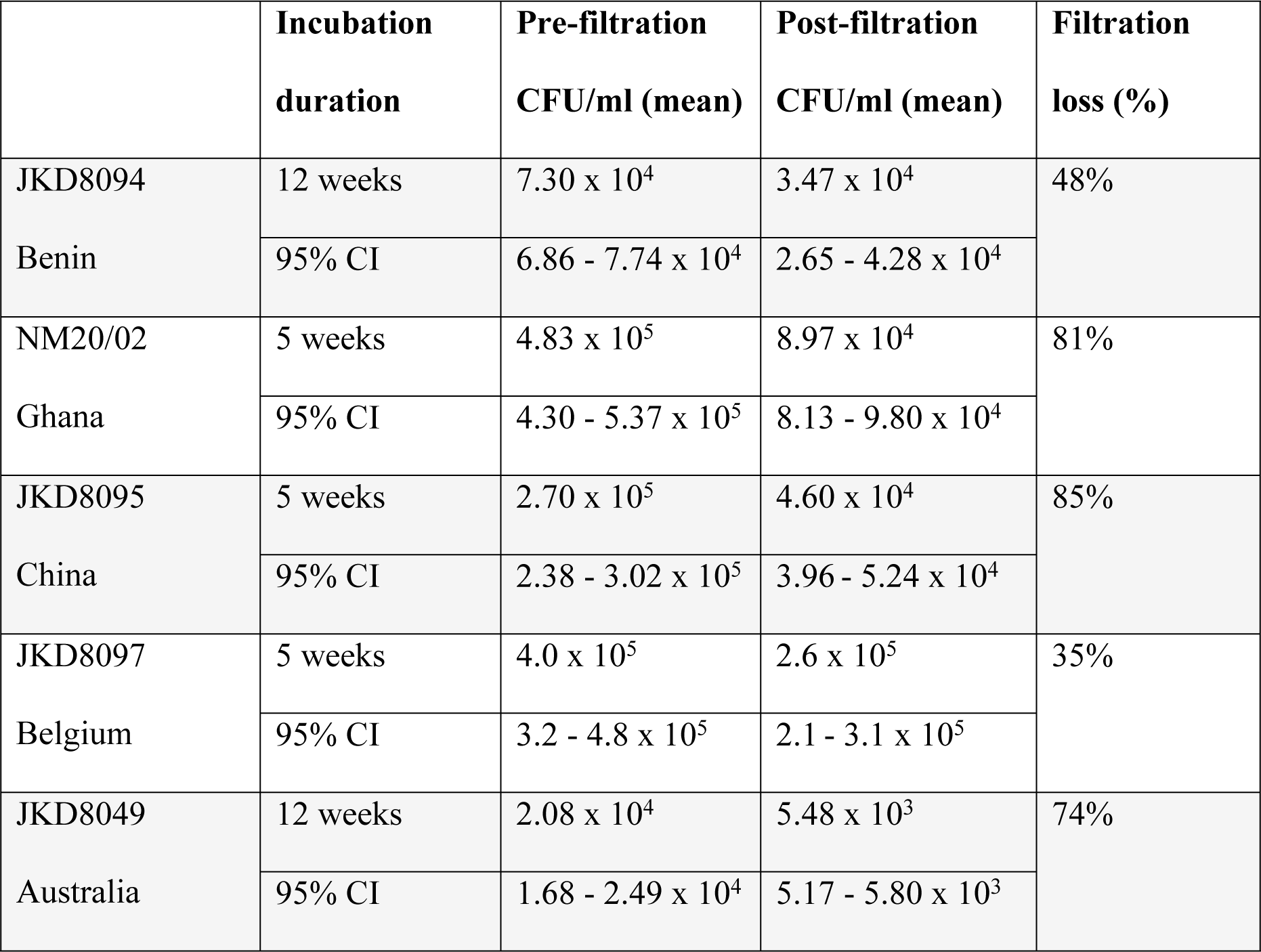
Mean CFU/mL and 95% confidence interval before and after filtration of orbital shaking cultures with glass beads.

**Table S3.** Mean CFU/mL, 95% confidence interval and viability loss compared to expected CFU/ml prior to glycerol storage in -80°C (accounting for dilution factor).

|  | **JKD 8094** | **NM20/02** | **JKD 8049** | **JKD 8095** | **JKD 8097** |
| --- | --- | --- | --- | --- | --- |
| **TTP:** | 10 weeks | 10 weeks | 6 weeks | 3 weeks | 2 weeks |
| **0 hr** | **2.17 x 10<sup>3</sup>** | **2.67 x 10<sup>4</sup>** | **1.83 x 10<sup>3</sup>** | **3.23 x 10<sup>3</sup>** | **2.43 x 10<sup>4</sup>** |
| 95% CI | 0.95 - 3.38 x10<sup>3</sup> | 1.81 - 3.52 x10<sup>4</sup> | 1.68 - 1.98 x10<sup>3</sup> | 2.83 - 3.64 x10<sup>3</sup> | 1.74 - 3.13 x10<sup>4</sup> |
| % loss | 81% | 11% | 0% | 79% | 72% |
| **1 hr** | **2.20 x 10<sup>3</sup>** | **2.37 x 10<sup>4</sup>** | Not tested | **3.40 x 10<sup>3</sup>** | **2.53 x 10<sup>4</sup>** |
| 95% CI | 1.27 - 3.13 x10<sup>3</sup> | 1.74 - 3.00 x10<sup>4</sup> |  | 2.84 - 3.96 x10<sup>3</sup> | 2.14 - 2.92 x 10<sup>4</sup> |
| % loss | 81% | 21% |  | 78% | 71% |
| **2 hrs** | **1.53 x 10<sup>3</sup>** | **3.00 x 10<sup>4</sup>** | **1.76 x 10<sup>3</sup>** | **3.77 x 10<sup>3</sup>** | Uncountable |
| 95% CI | 0.90 - 2.17 x10<sup>3</sup> | 2.56 - 3.44 x10<sup>4</sup> | 1.66 - 1.85 x10<sup>3</sup> | 3.21 - 4.32 x 10<sup>3</sup> |  |
| % loss | 87% | 0% | 4% | 75% |  |
| **4 hrs** | Not tested | Not tested | **1.66 x10<sup>3</sup>** | Not tested | Not tested |
| 95% CI |  |  | 1.54 - 1.78 x 10<sup>3</sup> |  |  |
| % loss |  |  | 9% |  |  |

TTP: Time-to-positive from thawing to final CFU enumeration; ‘% loss’ is the percentage reduction in viable cell count compared to prior to cryopreservation, accounting for dilution in glycerol storage media; 95% CI: 95% Confidence Interval.

**Fig. S1:**
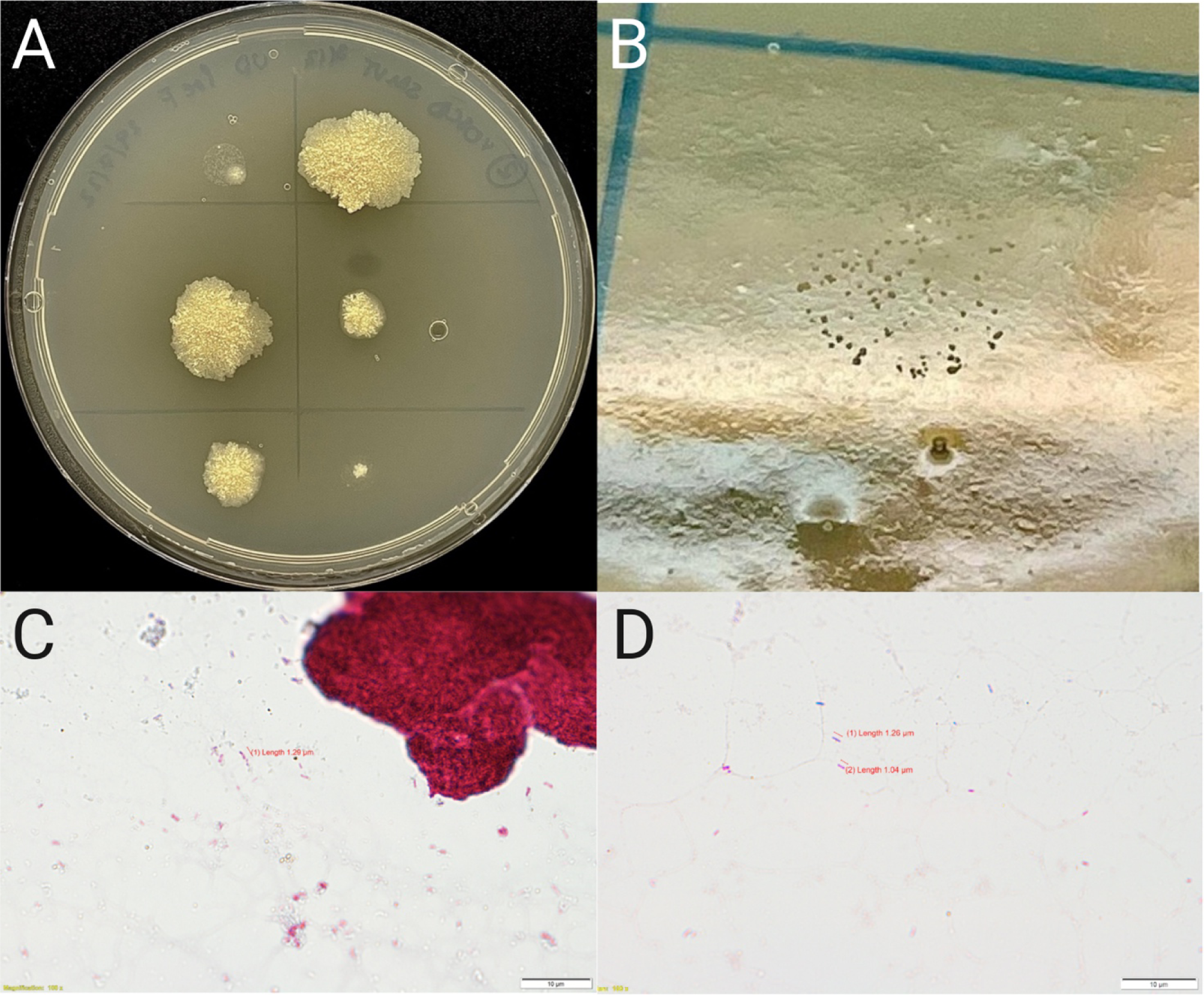
Pre-filtration (A) and post-filtration (B) spot plates of *M. ulcerans* JKD8049; pre-filtration spots demonstrate irregular colonies, due to the presence of clumps, which quickly overgrow other colonies and make enumeration difficult. Post-filtration spots demonstrate separated, countable microcolonies. Ziehl-Neilson stain of *M. ulcerans* JKD8049 cultured in orbital shaker with glass beads for 12 weeks before (C) filtration, at 100x magnification; occasional large clumps are visible, with a background of numerous individual bacilli; (D) Ziehl-Neilson stain of *M. ulcerans* JKD8049 filtered through a 5 μm pore filter, at 100x magnification. Bacilli are 1.0 to 1.5 μm in length, with no clumps visible in > 30 high power fields.

**Fig. S2.**
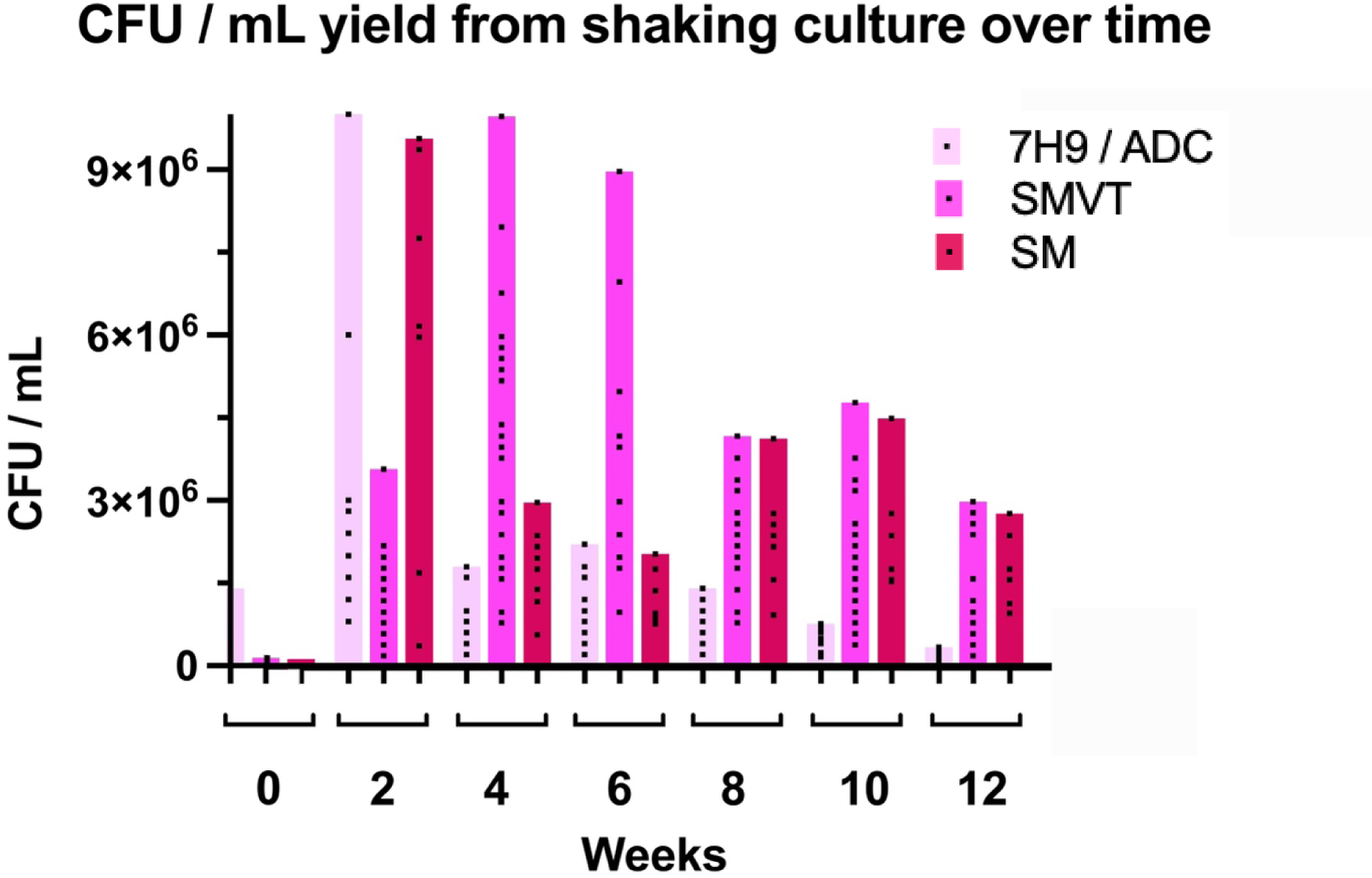
JKD8049 CFU/mL harvested following shaking culture in three different media; 7H9/ADC broth supplemented with albumin, dextrose and catalase (ADC), Sauton’s media (SM) with Veggietones (SMVT) and SM without nutritional supplementation. Samples were taken from the centre of the shaking culture, after pausing shaking for 20 minutes. Square boxes represent the CFU/mL obtained from each technical replicate. Six biological replicates were performed using SMVT, and three biological replicates for SM and 7H9/ADC, respectively.

## References

1. Muhi S, Osowicki J, O’Brien D, Johnson PDR, Pidot S, Doerflinger M, et al. A human model of Buruli ulcer: The case for controlled human infection and considerations for selecting a *Mycobacterium ulcerans* challenge strain. PLOS Neglected Trop Dis. 2023;17(6):e0011394.

2. Wallace JR, Mangas KM, Porter JL, Marcsisin R, Pidot SJ, Howden B, et al. *Mycobacterium ulcerans* low infectious dose and mechanical transmission support insect bites and puncturing injuries in the spread of Buruli ulcer. Plos Neglect Trop D. 2017;11(4):e0005553.

3. Röltgen K, Stinear TP, Pluschke G. The genome, evolution and diversity of *Mycobacterium ulcerans*. Infect Genetics Evol. 2012;12(3):522–9.

4. Oliveira MS, Fraga AG, Torrado E, Castro AG, Pereira JP, Filho AL, et al. Infection with *Mycobacterium ulcerans* induces persistent inflammatory responses in mice. Infect Immun. 2005;73(10):6299–310.

5. Bénard A, Sala C, Pluschke G. *Mycobacterium ulcerans* mouse model refinement for pre-clinical profiling of vaccine candidates. Plos One. 2016;11(11):e0167059.

6. Omansen TF, Porter JL, Johnson PDR, Werf TS van der, Stienstra Y, Stinear TP. *In-vitro* activity of avermectins against *Mycobacterium ulcerans*. Plos Neglect Trop D. 2015;9(3):e0003549.

7. Fyfe JAM, Lavender CJ, Handasyde KA, Legione AR, O’Brien CR, Stinear TP, et al. A major role for mammals in the ecology of *Mycobacterium ulcerans*. Plos Neglect Trop D. 2010;4(8):e791.

8. Faber WR, Arias-Bouda LMP, Zeegelaar JE, Kolk AHJ, Fonteyne PA, Toonstra J, et al. First reported case of *Mycobacterium ulcerans* infection in a patient from China. T Roy Soc Trop Med H. 2000;94(3):277–9.

9. Ranger BS, Mahrous EA, Mosi L, Adusumilli S, Lee RE, Colorni A, et al. Globally distributed mycobacterial fish pathogens produce a novel plasmid-encoded toxic macrolide, mycolactone F. Infect Immun. 2006;74(11):6037–45.

10. Price MN, Dehal PS, Arkin AP. FastTree: Computing large minimum evolution trees with profiles instead of a distance matrix. Mol Biol Evol. 2009;26(7):1641–50.

11. Siddiqi SH, Rüsch-Gerdes S. MGIT Procedure Manual. 2006.

12. Li G, Lian L lu, Wan L, Zhang J, Zhao X, Jiang Y, et al. Antimicrobial susceptibility of standard strains of nontuberculous mycobacteria by microplate alamar blue assay. Plos One. 2013;8(12):e84065.

13. Hoffner SE, Klintz L, Olsson-Liljequist B, Bolmström A. Evaluation of Etest for rapid susceptibility testing of *Mycobacterium chelonae* and *M. fortuitum*. J Clin Microbiol. 1994;32(8):1846–9.

14. Brown BA, Wallace RJ, Onyi GO, Rosas VD, Wallace RJ. Activities of four macrolides, including clarithromycin, against *Mycobacterium fortuitum*, Mycobacterium chelonae, and M. chelonae-like organisms. Antimicrob Agents Ch. 1992;36(1):180–4.

15. Biehle JR, Cavalieri SJ, Saubolle MA, Getsinger LJ. Evaluation of Etest for susceptibility testing of rapidly growing mycobacteria. J Clin Microbiol. 1995;33(7):1760–4.

16. Parish T, Stoker NG, editors. Mycobacteria protocols. Totowa, NJ: Humana Press; 1998.

17. Cheng N, Porter MA, Frick LW, Nguyen Y, Hayden JD, Young EF, et al. Filtration improves the performance of a high-throughput screen for anti-mycobacterial compounds. Plos One. 2014;9(5):e96348.

18. Fyfe JAM, Lavender CJ, Johnson PDR, Globan M, Sievers A, Azuolas J, et al. Development and application of two multiplex real-time PCR assays for the detection of *Mycobacterium ulcerans* in clinical and environmental samples. Appl Environ Microbiol. 2007;73(15):4733–40.

19. Stutz MD, Ojaimi S, Allison C, Preston S, Arandjelovic P, Hildebrand JM, et al. Necroptotic signaling is primed in *Mycobacterium tuberculosis*-infected macrophages, but its pathophysiological consequence in disease is restricted. Cell Death Differ. 2018;25(5):951–65.

20. Phillips RO, Robert J, Abass KM, Thompson W, Sarfo FS, Wilson T, et al. Rifampicin and clarithromycin (extended release) versus rifampicin and streptomycin for limited Buruli ulcer lesions: a randomised, open-label, non-inferiority phase 3 trial. Lancet. 2020;395(10232):1259–67.

21. O’Brien DP, Jenkin G, Buntine J, Steffen CM, McDonald A, Horne S, et al. Treatment and prevention of *Mycobacterium ulcerans* infection (Buruli ulcer) in Australia: guideline update. Med J Australia. 2014;200(5):267–70.

22. Venkataswamy MM, Goldberg MF, Baena A, Chan J, Jacobs WR, Porcelli SA. *In vitro* culture medium influences the vaccine efficacy of *Mycobacterium bovis* BCG. Vaccine. 2012;30(6):1038– 49.

23. Mve-Obiang A, Remacle J, Palomino JC, Houbion A, Portaels F. Growth and cytotoxic activity by *Mycobacterium ulcerans* in protein-free media. Fems Microbiol Lett. 1999;181(1):153–7.

24. Osowicki J, Azzopardi KI, McIntyre L, Rivera-Hernandez T, Ong C lynn Y, Baker C, et al. A controlled human infection model of group A streptococcus pharyngitis: Which strain and why? Msphere. 2019;4(1):e00647–18.

25. Malyala P, Singh M. Endotoxin limits in formulations for preclinical research. J Pharm Sci. 2008;97(6):2041–4.

26. Mve-Obiang A, Lee RE, Portaels F, Small PLC. Heterogeneity of mycolactones produced by clinical isolates of *Mycobacterium ulcerans*: Implications for virulence. Infect Immun. 2003;71(2):774–83.

27. Scherr N, Gersbach P, Dangy JP, Bomio C, Li J, Altmann KH, et al. Structure-activity relationship studies on the macrolide exotoxin mycolactone of *Mycobacterium ulcerans*. Plos Neglect Trop D. 2013;7(3):e2143.

28. Eddyani M, Portaels F. Survival of *Mycobacterium ulcerans* at 37°C. Clin Microbiol Infec. 2007;13(10):1033–5.

29. Zingue D, Panda A, Drancourt M. A protocol for culturing environmental strains of the Buruli ulcer agent, Mycobacterium ulcerans. Sci Rep-uk. 2018;8(1):6778.

30. Muhi S, Stinear TP. Systematic review of *M. bovis* BCG and other candidate vaccines for Buruli ulcer prophylaxis. Vaccine. 2021;39(50):7238–52.

31. Ishwarlall TZ, Adeleke VT, Maharaj L, Okpeku M, Adeniyi AA, Adeleke MA. Identification of potential candidate vaccines against *Mycobacterium ulcerans* based on the major facilitator superfamily transporter protein. Front Immunol. 2022;13:1023558.

32. Harris SA, Meyer J, Satti I, Marsay L, Poulton ID, Tanner R, et al. Evaluation of a human BCG challenge model to assess antimycobacterial immunity induced by BCG and a candidate tuberculosis vaccine, MVA85A, alone and in combination. J Infect Dis. 2014;209(8):1259–68.

33. Clinical and Laboratory Standards Institute (CLSI). Performance Standards for Susceptibility Testing of Mycobacteria, Nocardia spp., and Other Aerobic Actinomycetes. 2nd edition. CLSI supplement M24S. Clinical and Laboratory Standards Institute; 2023.

34. Aubry A, Jarlier V, Escolano S, Truffot-Pernot C, Cambau E. Antibiotic susceptibility pattern of *Mycobacterium marinum*. Antimicrob Agents Ch. 2000;44(11):3133–6.

35. Owusu E, Newman MJ, Addo KK, Addo P. In vitro susceptibility of *Mycobacterium ulcerans* isolates to selected antimicrobials. Can J Infect Dis Medical Microbiol J Can Des Maladies Infect Et De La Microbiol Médicale. 2017;2017:5180984.

36. Tobias NJ, Doig KD, Medema MH, Chen H, Haring V, Moore R, et al. Complete genome sequence of the frog pathogen *Mycobacterium ulcerans* ecovar Liflandii. J Bacteriol. 2013;195(3):556–64.

37. Pidot SJ, Porter JL, Lister T, Stinear TP. *In vitro* activity of SPR719 against *Mycobacterium ulcerans*, Mycobacterium marinum and Mycobacterium chimaera. Plos Neglect Trop D. 2021;15(7):e0009636.

38. Ji B, Lefrançois S, Robert J, Chauffour A, Truffot C, Jarlier V. *In vitro* and *in vivo* activities of rifampin, streptomycin, amikacin, moxifloxacin, R207910, linezolid, and PA-824 against *Mycobacterium ulcerans*. Antimicrob Agents Ch. 2006;50(6):1921–6.

39. Portaels F, Traore H, Ridder KD, Meyers WM. *In vitro* susceptibility of *Mycobacterium ulcerans* to clarithromycin. Antimicrob Agents Ch. 1998;42(8):2070–3.

40. Chu SY, Sennello LT, Sonders RC. Simultaneous determination of clarithromycin and 14(R)-hydroxyclarithromycin in plasma and urine using high-performance liquid chromatography with electrochemical detection. J Chromatogr B Biomed Sci Appl. 1991;571(1–2):199–208.

41. Periti P, Mazzei T. Clarithromycin: Pharmacokinetic and pharmacodynamic interrelationships and dosage regimen. J Chemotherapy. 2013;11(1):11–27.

42. Fraschini F, Scaglione F, Demartini G. Clarithromycin clinical pharmacokinetics. Clin Pharmacokinet. 1993;25(3):189–204.

43. Ruf MT, Steffen C, Bolz M, Schmid P, Pluschke G. Infiltrating leukocytes surround early Buruli ulcer lesions, but are unable to reach the mycolactone producing mycobacteria. Virulence. 2017;8(8):1918–26.

44. Torrado E, Fraga AG, Castro AG, Stragier P, Meyers WM, Portaels F, et al. Evidence for an intramacrophage growth phase of *Mycobacterium ulcerans*. Infect Immun. 2007;75(2):977–87.

45. Mitchison DA. Drug resistance in tuberculosis. Eur Respir J. 2005;25(2):376–9.

46. Yu X, Jiang G, Li H, Zhao Y, Zhang H, Zhao L, et al. Rifampin stability in 7H9 broth and Löwenstein-Jensen medium. J Clin Microbiol. 2011;49(3):784–9.

47. Dhungel L, Benbow ME, Jordan HR. Linking the *Mycobacterium ulcerans* environment to Buruli ulcer disease: Progress and challenges. One Heal. 2021;13:100311.

48. Buultjens AH, Vandelannoote K, Meehan CJ, Eddyani M, Jong BC de, Fyfe JAM, et al. Comparative genomics shows that *Mycobacterium ulcerans* migration and expansion preceded the rise of Buruli ulcer in southeastern Australia. Appl Environ Microb. 2018;84(8):e02612–17.

